# β-Catenin associates with a Wnt signaling mRNA network in myeloid cells through canonical RBP binding

**DOI:** 10.1101/2024.01.28.577638

**Authors:** M Wagstaff, O Sevim, A Goff, M Raynor, H Park, E Mancini, DTT Nguyen, T Chevassut, A Blair, L Castellano, S Newbury, B Towler, RG Morgan

## Abstract

Wnt/β-catenin signaling is important for normal hematopoietic stem/progenitor cell (HSPC) biology and heavily implicated in acute myeloid leukaemia (AML). The central mediator β-catenin is an attractive therapeutic target in AML however its targeting has been hampered by poor characterisation of its molecular interactions in haematopoietic cells. Our previous β-catenin interactome study identified the significant enrichment of RNA-binding proteins (RBP) implying post-transcriptional roles for β-catenin in myeloid cells. To identify β-catenin-associated mRNAs we performed β-catenin RNA-immunoprecipitation coupled to RNA-sequencing (RIP-seq) and identified significantly enriched Wnt signalling pathway transcripts. Using β-catenin cross-linking immunoprecipitation (CLIP) we demonstrated a limited capacity for β-catenin to bind RNA directly implying dependence on other RBPs. β-Catenin was found to interact with MSI2 in both myeloid cell lines and AML patient samples, where expression was significantly correlated. MSI2 knockdown reduced Wnt signalling output (TCF/LEF activity), through suppression of LEF-1 expression and nuclear localization. Through both RIP and CLIP we demonstrate MSI2 binds *LEF1* mRNA in a partly β-catenin dependent fashion, and may impact the post-transcriptional control of LEF-1 expression. Finally, we show that MSI2-mediated expansion of human HSPCs could be partly driven through *LEF1* regulation. This is the first study to show functional crosstalk between MSI2 and Wnt signalling in human cells, and indicates potential novel post-transcriptional roles for β-catenin in a haematological context.

## Introduction

Canonical Wnt signalling is an evolutionarily conserved pathway, critical in normal development and disease.^1^ In normal haematopoiesis, the central mediator β-catenin regulates the self-renewal^2^ and differentiation^3–5^ of haematopoietic stem/progenitor cells (HSPC), where its expression is tightly regulated to ensure the correct balance between HSPC maintenance and development.^6,7^ However its dysregulation is observed across numerous hematological malignancies, including myeloid leukaemias. Overexpression of β-catenin occurs in up to 80% of acute myeloid leukaemia (AML) cases,^8^ through a multitude of mechanisms,^9^ that result in hyperactive Wnt signalling^10^ and adverse patient survival.^11^ Deregulated Wnt signalling contributes to the development of several AML subtypes including normal karyotype,^12^ FLT3^-^mutant,^13^ del(5q)^+^,^14^ myelodysplastic syndrome (MDS)-related,^15^ CBF mutated,^16^ and PML-RARα^+^.^17^ Mouse models have also shown β-catenin regulates the emergence, maintenance and drug resistance of leukaemia stem cells (LSC) in both AML^18–23^ and chronic myeloid leukaemia (CML).^24,25^ This, coupled with the observation that only very low levels of Wnt/β-catenin signalling sustain normal haematopoiesis,^6^ makes the pathway an attractive therapeutic target. Yet despite this promise, pharmacological attempts to disrupt β-catenin have achieved limited success to date, hampered by a poor understanding of its molecular interactions in a haematopoietic context.

In solid tissues β-catenin has important cell adhesion functions through interaction with cadherins in adherens junctions.^26^ However, such tight cell-cell adhesion is not prominent in a fluid tissue like blood, meaning free β-catenin likely forms alternative interactions in haematopoietic cells. Indeed, our previous evaluation of β-catenin localisation in myeloid cells has not indicated significant plasma cell membrane localisation.^27^ To address this knowledge gap and identify new approaches for targeting β-catenin in haematological malignancy, we characterised β-catenin’s interaction network in myeloid cells and discovered a plethora of novel protein partners.^8^ Of note was the significant enrichment of RNA-binding proteins (RBP).^9^ RBPs are a diverse family of proteins that bind and regulate the stability, splicing, transport and translation of target RNAs, and are heavily implicated in normal hematopoiesis and leukaemia.^28^ Such interactions imply that β-catenin could have a hitherto uncharacterised but important role in post-transcriptional gene expression in haematopoietic cells.

In this study we show for the first time that β-catenin associates with a Wnt signalling mRNA network mediated through interactions with canonical RBPs like MSI2. We demonstrate MSI2 interacts with β-catenin and influences Wnt signaling output through modulation of the Wnt transcription factor LEF-1, which could be an important axis for controlling the growth and survival of human HSPCs.

## Methods

### Primary samples

Bone marrow, peripheral blood or leukapheresis samples from patients diagnosed with AML (Clinical information in Supplementary Table S1) were collected in accordance with the Declaration of Helsinki and with approval of University Hospitals Bristol NHS Trust and London Brent Research Ethics Committee. Human cord blood (CB) was obtained following informed consent healthy mothers at full-term undergoing elective caesarian sections at the Royal Sussex County Hospital, with approval from Brighton & Sussex University Hospitals NHS (BSUH) trust, the East of England – Essex Research Ethics Committee and HRA and Health and Care Research Wales (18/EE/0403). From all primary samples mononuclear cells (MNC) were isolated via density gradient separation using Ficoll-Hypaque (Merck Millipore) and only samples with ≥80% viability included in the study. The CD34^+^ fraction was derived as previously described^29^ from cryopreserved cord blood MNC preparations and enriched to >85% purity using MiniMACS (Miltenyi Biotec) according to the manufacturer’s instructions.

### Cell culture and drug treatments

The myeloid cell lines K562, HL60, HEL, U937, PLB-985, NOMO1, OCI-AML3, EOLI, ML-1, THP-1, KU812 (The European Collection of Authenticated Cell Cultures) and OCI-AML2, MV4-11, KG-1, KG1a SET2, NB4 and MONOMAC6 (Leibniz Institute DSMZ-German Collection of Microorganisms and Cell Cultures GmbH) were confirmed mycoplasma-free and authenticated via short-tandem repeat (STR) analysis prior to general culture as previously.^30^ β-Catenin was stabilised using the GSK-3β inhibitor CHIR99021 (Merck-Millipore) as previously described,^8^ whilst transcription was inhibited via 2µg/ml Actinomycin D (Merck-Millipore) at the stated timepoints. Purified human CB CD34^+^ HSPC were isolated and cultured at 5×10^5^/mL in StemSpan SFEMII (Stemcell Technologies) supplemented with human stem cell factor (150ng/mL), human FLT3-ligand (150ng/mL), and human thrombopoietin (TPO; 20ng/mL; Proteintech Group).

### RNA Immunoprecipitation (RIP)

RIP analyses were performed using the MagnaRIP kit (Merck-Millipore) according to manufacturer’s guidelines. Briefly, 2×10^6^ cells were treated overnight with CHIR99021 or DMSO at 37°C with 5% CO_2_. Cells were collected and washed in PBS followed by resuspension in lysis buffer, incubation on ice and overnight storage −80°C. Antibody-bead complexes were prepared following manufacturers guidelines with 5μg mouse IgG (Becton Dickinson), rabbit IgG (Merck Millipore), β-catenin (Clone 14, Becton Dickinson), HuR (Clone 3A2, Invitrogen), MSI2 (EP1305Y, Abcam) or LIN28B (Cell signalling Technology) and incubated at room temperature for 30 minutes with rotation. The RIP lysates were thawed and incubated with antibody-bead complexes with rotation at 4°C overnight followed by washing of unbound antibody.

Each immunoprecipitation was resuspended in proteinase K buffer to remove contaminating protein and followed by addition of phenol: chloroform: isoamyl alcohol in a 25:24:1 (Thermofisher Scientific) to separate phases. Aqueous phase was removed to a new eppendorf followed by ethanol addition to precipitate RNA at - 80°C overnight. All tubes were centrifuged, followed by washing in 80% ethanol and then air-drying of pellet for 10 minutes at room temperature after a final centrifugation. The RNA pellet was resuspended in 20μl of nuclease-free water and the tubes kept at −80°C ahead of sample assessment by an mRNA bioanalyzer Pico Series II (Agilent).

### RIP sequencing analysis

Raw FASTQ files were trimmed using Cutadapt (version 4.6) to remove adapters identified during the quality control process.^31^ The trimmed reads were then mapped to the human genome using HISAT2 (version 2.2.1) with default parameters.^32^ The <u>human genome index file</u> used for alignment was downloaded from the HISAT2 website. SAMtools (version 1.21) was used to convert SAM files generated by HISAT2 into BAM files for read counting.^33^ Reads were then counted using featureCounts (version 2.0.6) with a human annotation file from <u>Ensembl</u>, applying the parameters: “--primary-C-t exon-g gene_id”. The raw read counts were imported into R to identify differentially enriched RNAs between basal (DMSO) and stabilised β-catenin (CHIR99021) conditions. Differential enrichment analysis, including fold change calculation and statistical analysis, was conducted with the DESeq2 package (version 1.44.0)^34^ in <u>R</u>. Data manipulation and visualization were performed using the tidyverse (version 2.0.0)^35^ and ComplexHeatmap (version 2.20.0)^36^ packages.

### Cross linking immunoprecipitation (CLIP)

4×10^7^ cells were harvested, washed twice with room-temperature PBS and split across four 60 mm x 15 mm cell culture dishes (Corning) prior to UV crosslinking. Crosslinking was performed using three rounds of 100mJ/cm² UV-254 nm irradiation, with agitation between each round. Following UV treatment, cells were recombined and centrifuged at before disposal of supernatant and cell pellets snap-frozen in liquid nitrogen. Lysates were resuspended in 500µL lysis buffer as previously,^27^ and sonicated for 10 cycles of alternate 30 seconds of sonication with 30-second rests.

Sonicated lysates were centrifuged to pellet debris followed by quantitation of lysates using the DC protein assay (BioRad) as previously described,^27^ to ensure equal protein input for downstream RT-qPCR. Four units of Turbo DNase (Ambion, Thermofisher) was added to each sample and incubated at 37°C for 3 minutes followed by cooling on ice for 3 minutes. 8µg β-catenin or matched murine IgG antibody (Becton Dickinson), MSI2 or LIN28B, and matched Rabbit IgG antibody (Cell Signalling Technology) was conjugated to beads as previously described,^27^ but using CLIP lysis buffer (50 mM Tris-HCl, pH 7.4, 100 mM NaCl, 1% Igepal CA-630, 1% sodium deoxycholate) without protease inhibitors. Lysates were incubated with antibody-bead complexes for a minimum of 1 hour at 4°C with rotation with 150µL original lysate retained for input samples, after which unbound antibody discarded from beads using washes in CLIP lysis buffer. Antibody-bead complexes were resuspended in 250µL Proteinase K elution buffer (50 mM Tris-HCl, pH 7.4, 100 mM NaCl, 1% Igepal CA-630, 1% sodium deoxycholate) with 20 mg/mL proteinase K solution (ThermoScientific) added to each sample, followed by incubation at 65°C for 1 hour prior to magnetic bead separation and supernatant transferred to fresh tubes. To each sample, 750µL TRIzol LS reagent (Invitrogen) was added before storage at −80°C for RNA extraction. RNA extraction, purification, and quantification were performed as above.

### RNA extraction, clean up and quantitation

RNA was extracted using the Zymo RNA mini-prep kit (Cambridge Bioscience), cleaned using RNeasy MinElute Clean-up kit (Qiagen), or by DNase treatment using the DNA free removal kit (Thermofisher), all via manufacturer guidelines.^37^ RNA was quantified using a Nanodrop 2000 spectrophotometer (Thermofisher) blanked against nuclease-free water.

### RT-qPCR

Purified RNA was converted to cDNA using the high-capacity RNA-to-cDNA kit (Thermofisher) or the one-step RT-qPCR mix (APTO-GEN, Raynham, Massachusetts, USA). All primers used in this study are listed in Supplementary Table S2 and were diluted to 10mM with qRT-PCR programmes used as per manufacturers guidelines (Merck-Millipore). RT-qPCR was performed on a QuantStudio™ 3 Real-Time PCR System (ThermoFisher Scientific) in conjunction with associated software. Relative gene expression was determined first by subtracting the C_T_ value of housekeeping genes (*ACTB*, *GAPDH* or *RNA18SN1*) from the C_T_ value of genes of interest to generate Δ^CT^. The ΔΔ^CT^ was then calculated by subtracting the average control Δ^CT^ from the Δ^CT^ of each condition and fold change calculated through 2^-ΔΔCT^.

To account for differences in RNA sample preparation, RIP RNA C_T_ values were normalised to the corresponding input RNA fraction C_T_ value for the same qPCR assay as per manufacturer’s guidelines. Briefly, Δ^CT^ values were calculated by subtracting the C_T_ value of the input fraction from the C_T_ value of the RIP fraction for each target gene, ensuring variability in RNA input or preparation was accounted for. In contrast, for CLIP analysis, normalisation of CLIP RNA to the input was not required, as sample concentrations were matched at the protein level prior to Co-IP.

### Co-immunoprecipitation (Co-IP)

Cross-linking and Co-immunoprecipitation was executed as previously.^8^ Briefly, 5μg of MSI2 (EP1305Y, Abcam, Oxford, UK), HuR (Clone 3A2, Invitrogen), β-catenin (Clone 14, Becton Dickinson), mouse IgG (Becton Dickinson) of rabbit IgG (Merck Millipore, New Jersey, United States) was complexed with Protein G Dynabeads (Invitrogen), and incubated with 1mg of whole cell lysate overnight in Co-IP lysis buffer (1 x Cell Signalling Technology lysis buffer, plus 10% glycerol and 0.5mM dithiothreitol) at 4°C or with the addition of 20μg/ml RNase A (Thermofisher Scientific). Following washing of antibody-bead complexes five times in Co-IP lysis buffer, each sample was diluted in 2x Laemmli buffer (ThermoFisher Scientific) plus Co-IP lysis buffer and heated to 95°C for 5 minutes prior to immunoblotting.

### Immunoblotting

Immunoblotting was performed as previously,^8^ using antibodies to β-catenin, MSI2, LEF-1, GAPDH (Proteintech, Manchester, UK), Lamin A/C (Merck Millipore, New Jersey, United States) and α-tubulin (Merck Millipore) already outlined in above sections.

### Lentivirus preparation

K562 and KU812 and cells were lentivirally transduced with the β-catenin-activated reporter (BAR) or mutant ‘found unresponsive’ control (fuBAR) system as previously.^8^ The lentiviral expression plasmids used for transgene expression in this study are listed in Supplementary Table S3 and were used alongside their appropriate matched controls.

### Flow cytometry

For Wnt reporter assessment, 2×10^5^ BAR/fuBAR containing cells were treated overnight with either DMSO or CHIR99021. TCF/LEF reporter activity (Venus YFP intensity) was evaluated by flow cytometric analysis using a CytoFLEX Flow cytometer (Beckman Coulter, California, USA) in conjunction with FlowJo software Version 10.10 (Tree Star Inc., Ashland, OR USA) acquiring a minimum of 2×10^4^ debris excluded events. For immunophenotyping, up to 5×10^4^ CB-derived CD34^+^ HSPC were seeded into wells of 96-well plate in 100µl staining buffer (1xPBS, 0.5% BSA) and stained with 10µg/mL antibodies to CD34-PE (Clone 581, Biolegend) and CD45-PerCPCy5.5 (Clone QA17A19, Biolegend) or the equivalent concentration-, manufacturer- and isotype-matched control antibodies (Clone MOPC-21, Biolegend).

### Nuclear/cytoplasmic fractionation

Nuclear/cytoplasmic fractionation was performed as previously.^8^

### Statistics

For densitometry, ImageJ Version 2.9.0 (National Institute of Health) was used to quantify regions of interest which was normalised to the GAPDH intensity obtained within each respective sample. Spearman Rank correlation was assessed using Prism Version 10.3.1 (GraphPad Software, LLC), alongside statistical tests including one-sample or students t-test, using a threshold of *p*<0.05. Unless otherwise stated all tests were performed in biological triplicates with data representing the mean ± 1 standard deviation (s.d).

## Results

### β-Catenin RIP-seq reveals enrichment with Wnt signalling mRNAs

Our previous β-catenin interactome study in myeloid cells revealed the significant enrichment of RBPs,^8,9^ indicating a putative novel role for β-catenin in post-transcriptional processes in haematological cells. To first confirm whether β-catenin associates with RNA in myeloid cells we performed β-catenin RNA-immunoprecipitation (RIP) and examined RNA concentration and size using a Bioanalyser system. We first confirmed efficient RIP using K562 cells and a canonical RBP target, HuR (*ELAVL1,* also found to interact with β-catenin; Supplemental Figure S1),^38^ with a known mRNA binding partner *ACTB* detected by RT-qPCR (Figure 1B). Using this assay we next successfully enriched β-catenin via RIP from K562 cells under basal (DMSO) or Wnt signalling stimulated (5μM GSK3β inhibitor CHIR99021) conditions (Figure 1C) and showed abundant RNA enrichment versus IgG-RIP (Figure 1D).

**Figure 1.**
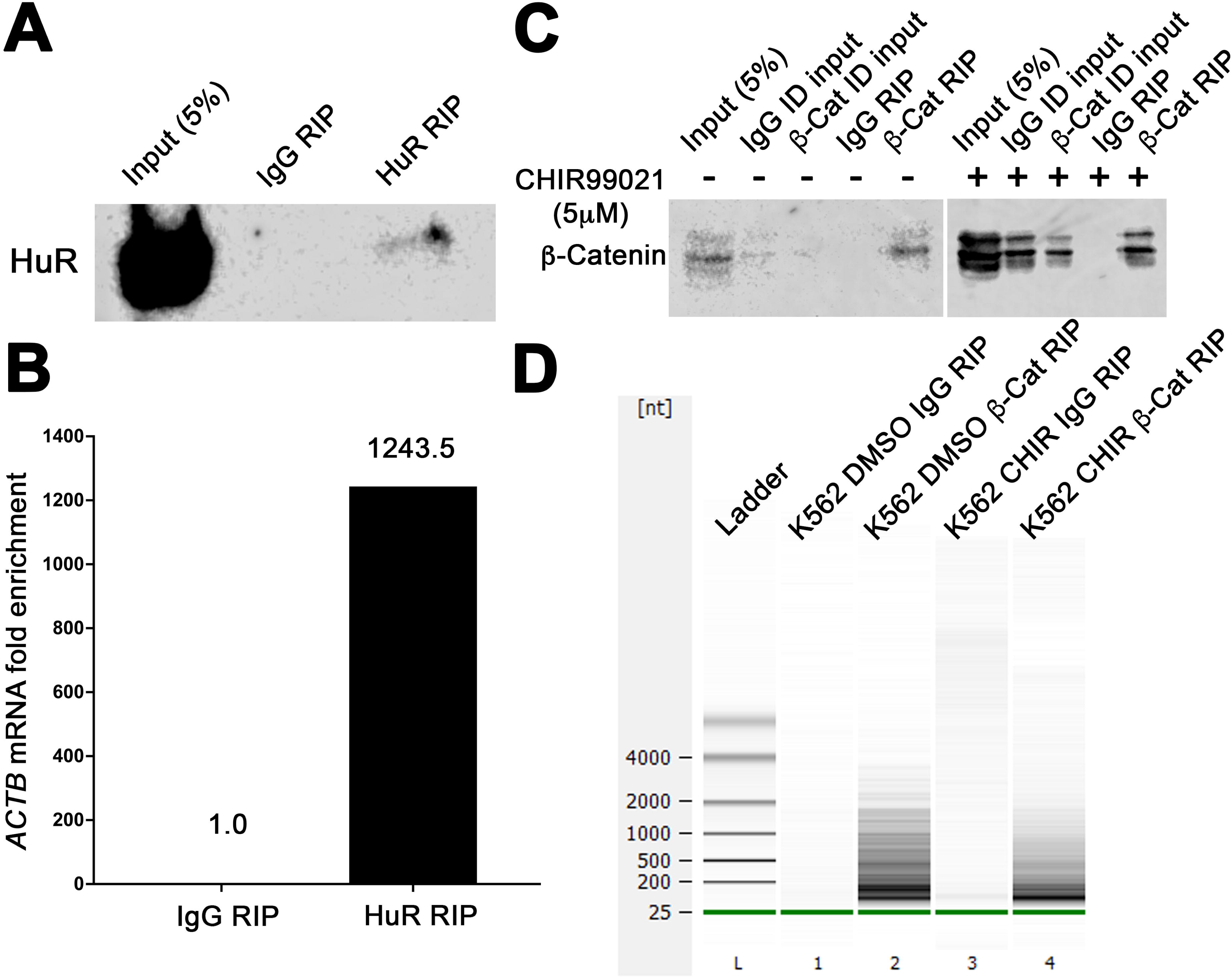
β-Catenin RIP enriches with RNA from myeloid cells. **A**) Immunoblot showing HuR levels in HuR RIP performed from K562 cells. **B**) Graph summarising the fold enrichment *ACTB* mRNA (known HuR binding target)^75^ obtained from RT-qPCR analysis of IgG or HuR RIP analysis from K562 cells (n=1) **C**) Representative immunoblot showing β-catenin level obtained from β-catenin RIPs performed from K562 cells (+/- CHIR99021), ID= immunodepleted lysate. **D**) Agilent 2100 Bioanalyzer gel showing RNA isolated from IgG or β-catenin RIP performed from K562 cells (+/- CHIR99021).

Following successful β-catenin RIP we next coupled this to RNA-sequencing to identify β-catenin-associated transcripts. Using K562 and HEL cells which we have previously shown to be acutely Wnt signalling responsive,^8^ we identified 582 and 1179 mRNAs respectively, that were enriched (FDR *p*<0.05) between basal and stabilized β-catenin conditions, which included Wnt signalling transcripts (Figure 2A; raw β-catenin RIP-seq data publicly available at ArrayExpress under reference E-MTAB-14675, and processed data available as Supplementary Data File 1). Further gene ontology pathway analysis highlighted Wnt/β-catenin signalling mRNAs as amongst the most significantly enriched from K562 β-catenin RIP-seq (Figure 2B), and a number of these were also observed in the β-catenin RIP-seq from HEL cells (Figure 2C). Using β-catenin RIP-RT-qPCR (versus IgG RIP-RT-qPCR) we validated enrichment of several Wnt signalling mRNAs in K562 and HEL cells including *AMER1*, *BCL9L*, *AXIN2* and *LEF1* (Figure 2D and E). Taken together, these data show that β-catenin associates with mRNA in myeloid cells which are significantly enriched for Wnt signalling components.

**Figure 2.**
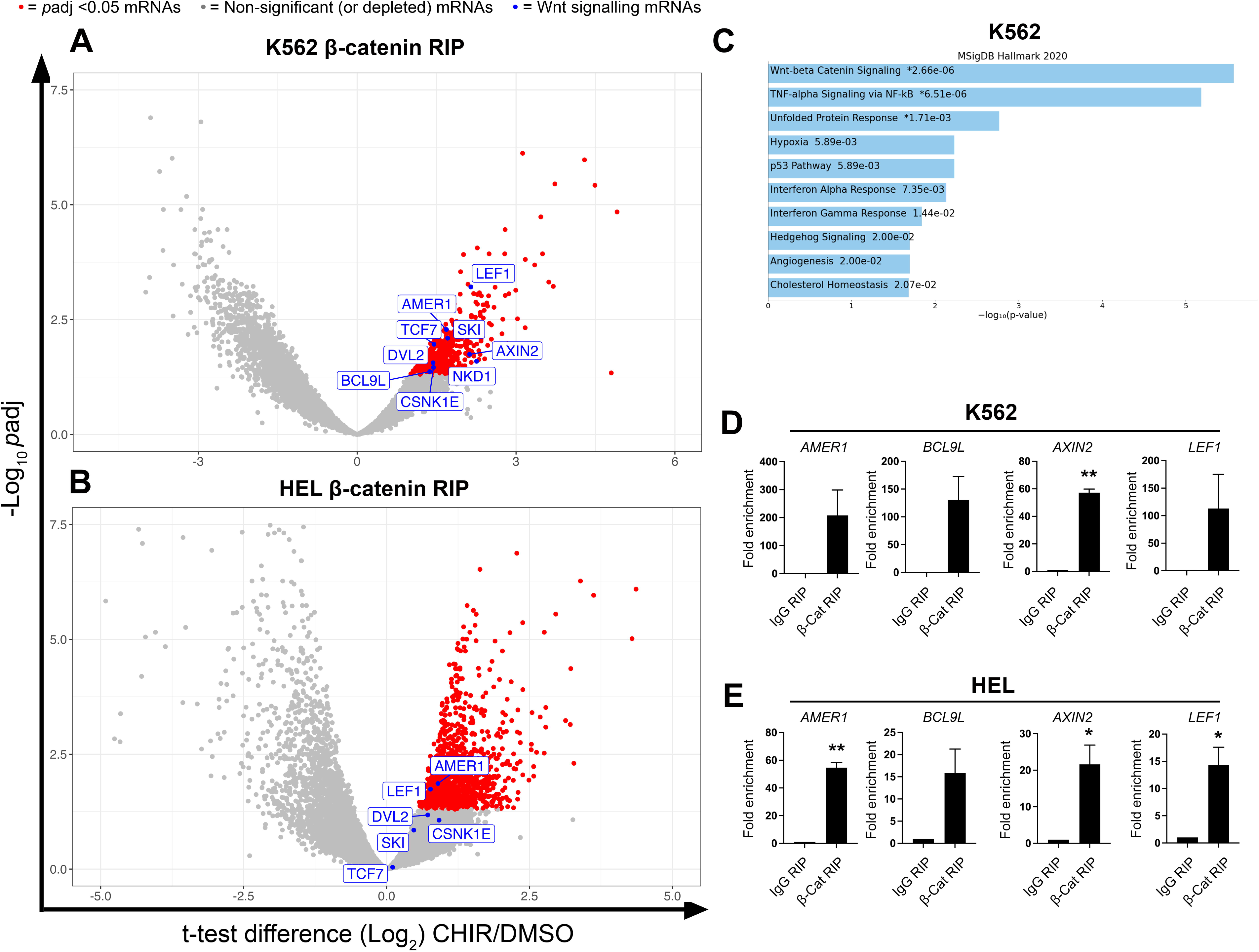
β-Catenin RIP enriches with Wnt signalling mRNAs. Volcano plots showing the fold change in mRNA abundance detected within β-catenin RIP performed from DMSO (basal Wnt signalling) versus CHIR99021 (activated Wnt signalling) treated **A**) K562 or **B**) HEL cells (n=3). Red dots represent enriched (*P*adj<0.05) mRNAs whilst blue dots highlight Wnt signalling mRNAs. **C**) Gene set enrichment analysis using the human Molecular Signatures Database (MSigDB Hallmark 2020) for pathways represented amongst the significantly enriched mRNAs (*P*adj<0.05), obtained in β-catenin RIP from K562 cells +/- CHIR99021 with adjusted-Log_10_ *P* values annotated. Summary graphs showing the fold enrichment of selected Wnt signalling mRNAs isolated from IgG or β-catenin RIP-RT-qPCR performed in **D**) K562 and **E**) HEL cells. Data represents mean ± 1 s.d (*n* = 3). Statistical analysis is denoted by \**p*<0.05 and \*\**p*<0.01 as deduced from a student’s t-test.

### β-Catenin has weak capacity for RNA binding in myeloid cells

β-Catenin lacks a canonical RNA-binding motif however intrinsically disordered regions (IDR), which comprise β-catenin’s N- and C-termini (Supplemental Figure S2),^39^ are commonly implicated in RNA binding by non-canonical RBPs.^40^ Furthermore, Armadillo domains are the one of the most frequently implicated structures in non-canonical RNA binding and β-catenin contains 12 such repeating domains in its central region which predict RNA binding (Supplemental Figure S3).^41^

β-Catenin has been reported to directly bind the 3’-UTR of mRNAs,^38,42^ and has recently been shown to bind double-stranded RNA.^43^ Therefore, to assess the RNA binding capacity of β-catenin in the haematopoietic system we performed β-catenin cross-linking immunoprecipitation (CLIP), a more stringent method of RBP:RNA capture. As shown in Figure 3A, β-catenin RIP once again pulled down with abundant RNA relative to IgG RIP, as did a positive canonical RBP control with LIN28B RIP. However, using a CLIP approach, despite efficient β-catenin isolation (Figure 3B), there was no observable RNA enrichment over the control IgG CLIP, whilst LIN28B CLIP still pulled down abundant RNA as expected, suggesting β-catenin does not directly bind RNA strongly in this context.

**Figure 3.**
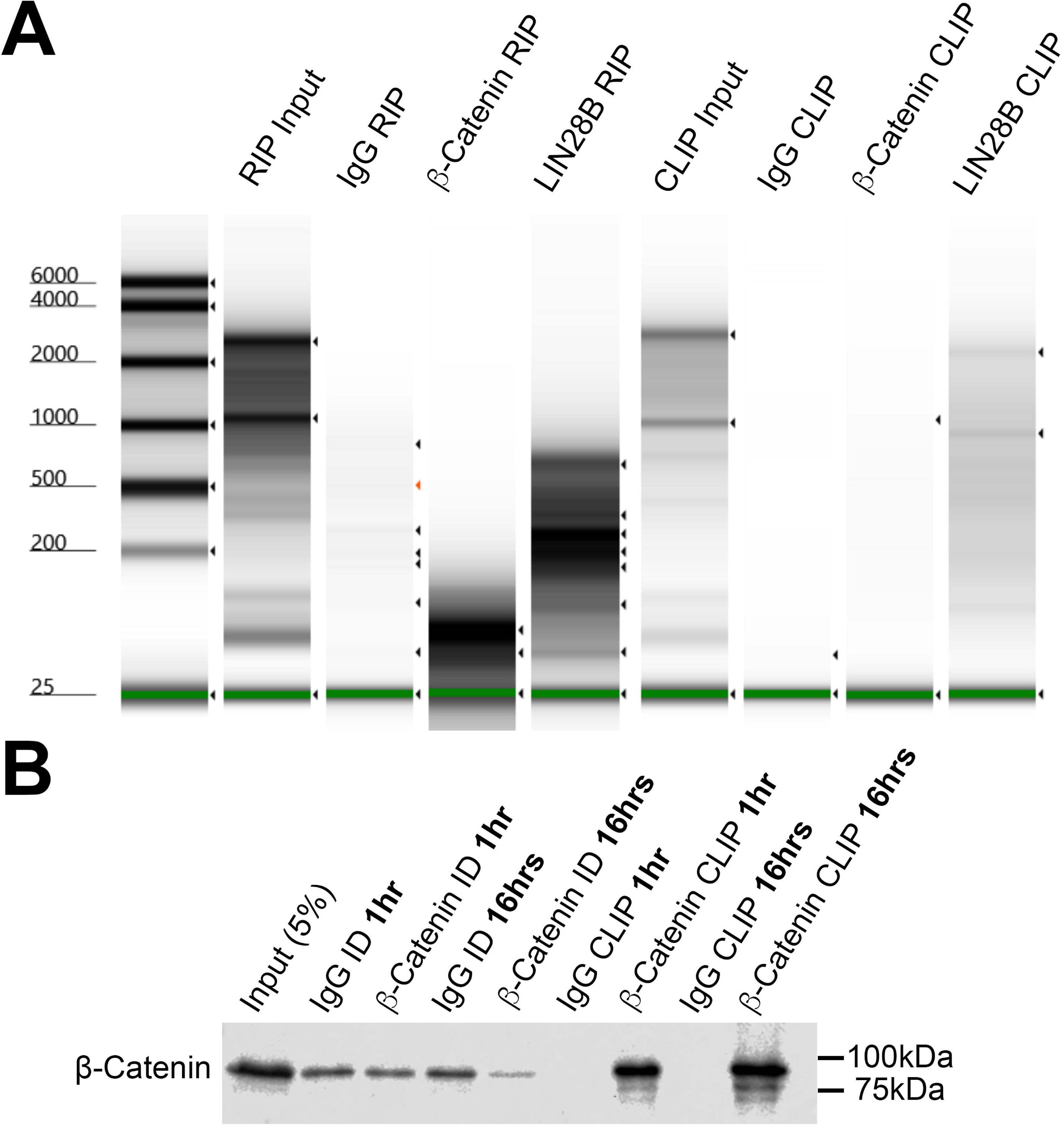
β-Catenin CLIP does not enrich with RNA from myeloid cells. **A**) Representative Agilent 2100 Bioanalyzer gel showing RNA isolated from IgG, β-catenin or LIN28B RIP and CLIP performed in K562 cells. **B**) Representative immunoblot showing β-catenin level in IgG or β-catenin RIP performed from K562 cells using 1hr or 16hr primary antibody incubation, ID= immunodepleted lysate.

### MSI2 interacts with β-catenin in myeloid cells

We anticipated that β-catenin might network with Wnt signalling mRNAs indirectly through other canonical RBPs and returned to our previously characterised β-catenin interaction network.^8,9^ Musashi-2 (MSI2) was identified as a putative β-catenin interacting RBP in myeloid cells (Figure 4A),^8^ and colorectal SW620 cells suggesting this interaction could also occur beyond haematopoietic tissue (Supplemental Figure S4). To confirm interaction, we performed reciprocal MSI2 co-immunoprecipitation (Co-IP) from K562 and HEL cells and confirmed consistent β-catenin enrichment under both basal and Wnt signalling activated conditions (Figure 4CD). Since MSI2 is an RBP,^44^ and β-catenin has also been shown to bind RNA,^38,42^ we assessed whether the β-catenin:MSI2 interaction could be a consequence of RNA co-occupancy. However, after repeating MSI2 co-IP +/-RNase A, and confirming complete digestion of RNA (Supplemental Figure S5),^27^ the MSI2:β-catenin protein interaction remained suggesting these proteins are complexed irrespective of RNA presence (Figure 5CD). Furthermore, using AlphaFold3 modelling, we found the accuracy of the predicted relative positions of the subunits in the β-catenin:MSI2 complex to be low, suggesting that direct interaction is unlikely and thus likely mediated by intermediary partners (Supplemental Figure 5).

**Figure 4.**
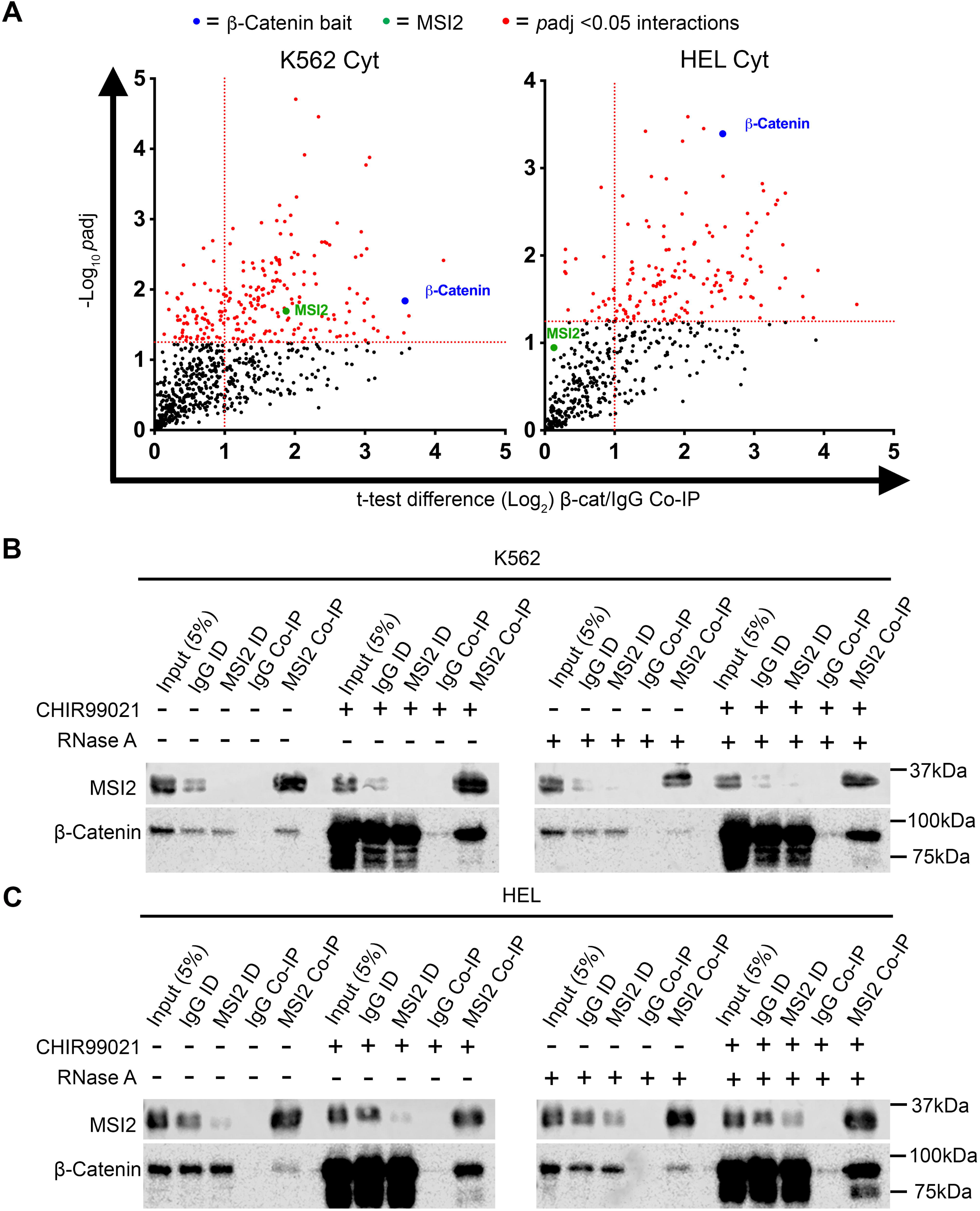
β-Catenin interacts with MSI2 in myeloid cells. **A**) Scatter plots showing MSI2 detection in β-catenin interactomes performed in cytosolic fractions of K562 and HEL cells. Vertical dashed red line indicates the threshold for 2-fold change in protein binding at log_2_ (=1) relative to IgG Co-IP. Horizontal red line represents threshold for *p*<0.05 on log_10_ scale (=1.3). Highlighted red dots indicate enriched interactions (*p*<0.05), green labels highlight position of MSI2, and blue labels highlight position of β-catenin bait. Representative immunoblots showing the level of β-catenin protein present in MSI2 Co-IPs derived from **B)** K562 and **C)** HEL whole cell lysates +/- 20µg/mL RNaseA +/-5µM CHIR99021 overnight. ID= immunodepleted lysate. Data represents mean ± 1 s.d (*n* = 3). Statistical analysis is denoted by \**p*<0.05, \*\**p*<0.01 and \*\*\*\**p*<0.0001 as deduced from a student’s t-test.

**Figure 5.**
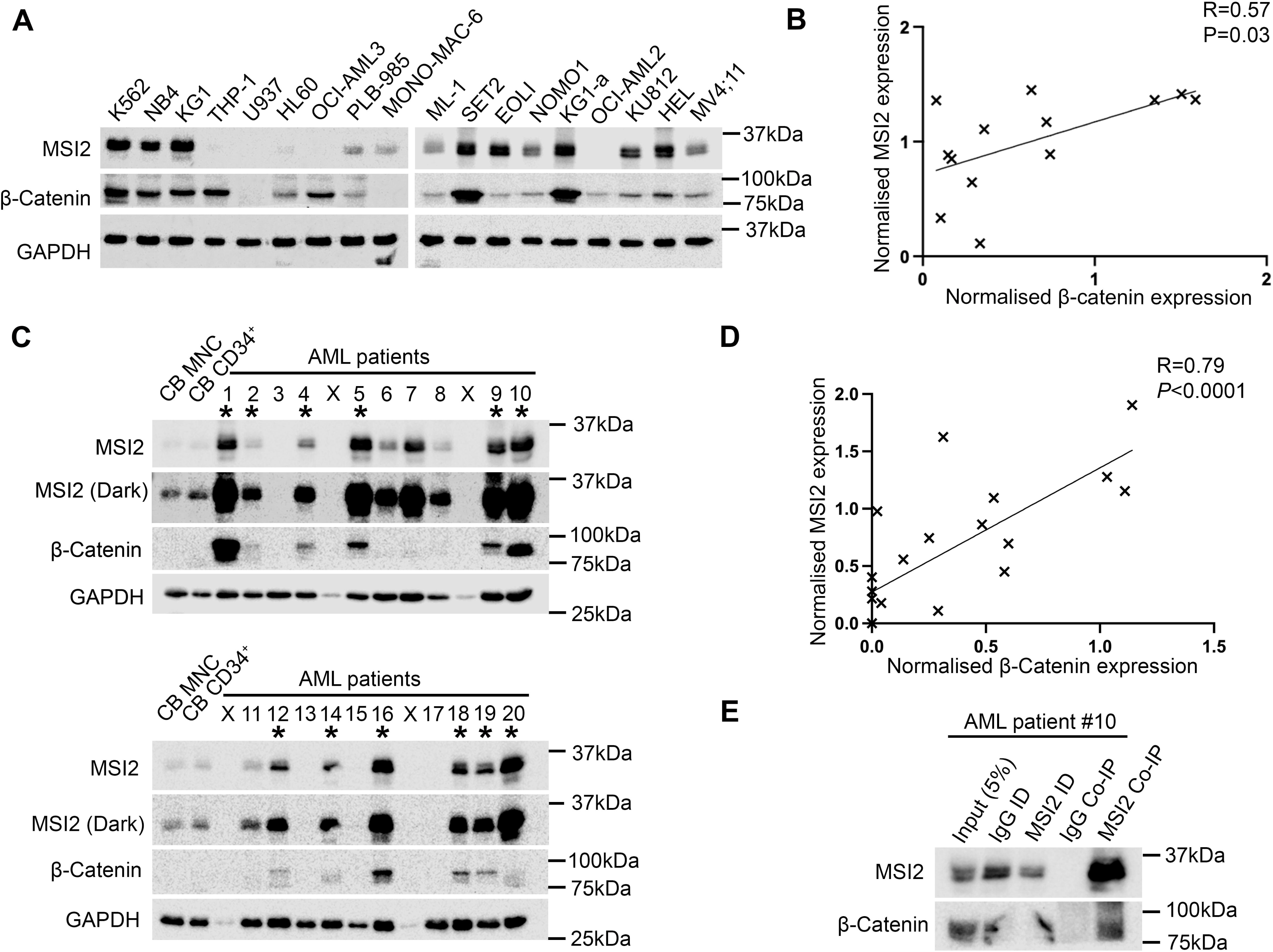
MSI2 and β-catenin correlate and interact in primary AML patient samples. **A)** Immunoblot of 18 myeloid leukemia cell lines showing the relative level of β-catenin and MSI2 protein, with GAPDH used to assess protein loading. **B)** Summary scatter plot showing the correlation (Spearman Rank R=0.6, P<0.03) between relative β-catenin and MSI2 protein expression across 18 myeloid cell lines (normalised to GAPDH expression within each cell line). **C)** Immunoblot screen of 20 primary AML patient samples showing the relative level of β-catenin and MSI2 protein (light and dark exposures). * denotes samples co-overexpressing both β-catenin and MSI2 relative to levels in CB MNC and CB CD34+ enriched fraction (pooled from five independent CB samples). X = Void sample containing no protein as deduced from negative GAPDH detection. **B)** Summary scatter plot showing the correlation (Spearman Rank R=0.79, P<0.0001) between relative β-catenin and MSI2 protein expression in 20 primary AML patient samples (normalised to GAPDH expression in each sample). **C)** Immunoblot showing the level of β-catenin protein present in an MSI2 Co-IP performed from primary AML patient sample #10 of sample screen.

### β-Catenin interacts with MSI2 in myeloid cell lines and primary AML cells

To assess the potential scope of cooperation between MSI2 and β-catenin across myeloid cells we assessed MSI2 expression across a panel of 18 myeloid cell lines. Two thirds of the myeloid cell lines assessed exhibited abundant co-expression of both β-catenin and MSI2 (Figure 5A) which exhibited a degree of correlation (*p*=0.03; Figure 5B). To explore the clinical relevance of the β-catenin:MSI2 association, we screened 20 AML patients for protein expression of MSI2 and β-catenin (patient clinical details provided in Supplemental Table S1). Like myeloid cell lines, MSI2 was frequently co-expressed with β-catenin to a similar frequency observed in myeloid cell lines (12/20 samples [60%]; Figure 5C), with an extensive correlation (*p*<0.001) between the two proteins (Figure 5D). To examine if the MSI2:β-catenin interaction could be detected in patient cells, we performed an MSI2 Co-IP in a primary AML sample containing abundant levels of both proteins and where ample cellular material existed (patient 10) and confirmed MSI2 interaction with β-catenin (Figure 5E). Taken together these data indicate the β-catenin:MSI2 interaction could represent an abundant and clinically relevant association in AML cells.

### MSI2 impacts Wnt signalling output through modulation of LEF-1

To evaluate if MSI2 could bind or regulate Wnt mRNAs we assessed its net impact on Wnt signalling output using the **b**eta-catenin **a**ctivated **r**eporter as done previously.^8,27,30^ Depletion of MSI2 using two unique shRNAs had little impact on β-catenin level but did cause a concomitant decrease in expression of the Wnt transcription factor LEF-1 (Figure 6A). As could be expected from diminished LEF-1, this resulted in diminished Wnt signalling output from both K562 and KU812 cells exhibiting MSI2 knockdown (Figure 6B-D). MSI2 knockdown in HEL cells also diminished Wnt signalling output and LEF-1 expression (Supplemental Figure S7), however we were unable to use this cell line for subsequent experiments since only one MSI2 shRNA variant remained viable after lentiviral transduction and selection. We also demonstrated that ectopic MSI2 expression increased nuclear LEF-1 expression and TCF/LEF activity in K562 cells (Supplemental Figure S8) in Wnt signalling activated conditions.

**Figure 6.**
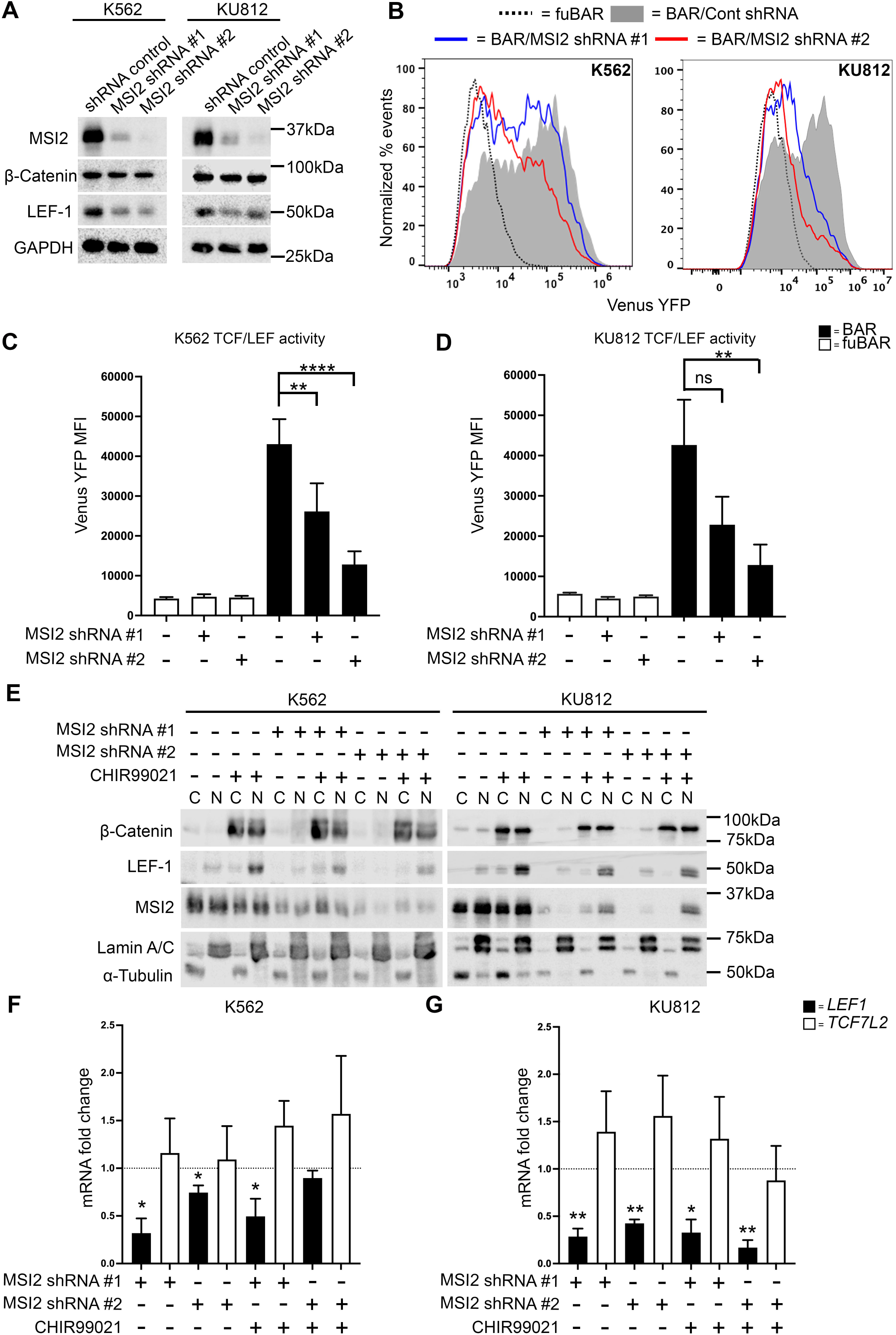
MSI2 knockdown impairs Wnt signalling output through LEF-1 modulation. **A)** Immunoblots showing MSI2, β-catenin and LEF1 level in K562 and KU812 cells harbouring MSI2 shRNA or non-targeting shRNA controls. GAPDH indicates protein loading. **B**) Representative flow cytometric histograms showing intensity of the TCF-dependent expression of Venus Yellow Fluorescent Protein (YFP) from the β-catenin activated reporter (BAR) reporter, or negative control ‘found unresponsive’ BAR (fuBAR; containing mutated promoter binding sites) in K562 and KU812 cells +/- MSI2 shRNA following treatment with 5μM CHIR99021 overnight. The fuBAR (dashed), non-targeting control shRNA (grey filled), and two MSI2 shRNAs (blue or red) histograms are shown. Summary graphs showing the median fluorescence intensity (MFI) generated from the BAR/fuBAR in **C**) K562 and **D**) KU812 cells +/- MSI2 shRNA with +/- 5μM CHIR99021. **E)** Immunoblots showing total β-catenin, LEF-1 and MSI2 subcellular localization in K562 and KU812 cells lentivirally transduced with two different MSI2 shRNAs +/- 5μM CHIR99021. Lamin A/C and α-tubulin indicate the purity/loading of the nuclear (N) and cytosol (C) fractions respectively. Summary graph showing the fold change in *LEF1* and *TCF7L2* mRNA expression as assessed by RT-qPCR in **F**) K562 and **G**) KU812 cells expressing MSI2 shRNA. Fold change is relative to matched respective controls (black dashed line) and overall expression was normalized to the housekeeping gene β-actin (*ACTB*). Data represents mean ± 1 s.d (*n* = 3). Statistical analysis is denoted by \**p*<0.05, \*\**p*<0.01 and \*\*\*\**p*<0.0001 as deduced from a student’s t-test.

To understand the basis for reduced TCF/LEF output in MSI2-deficient cells, we performed nuclear/cytoplasmic fractionation and examined the expression/localisation of Wnt signalling components. Again, we observed limited impact of MSI2 knockdown on β-catenin level or nuclear localisation in response to Wnt signalling activation but did note markedly reduced LEF-1 expression and nuclear localisation in K562 and KU812 cells under both basal and Wnt signalling active states (Figure 6E). To understand the basis for reduced LEF-1 expression upon MSI2 knockdown we examined mRNA levels using qRT-PCR and observed significantly reduced *LEF1* (but not that of another critical Wnt effector; *TCF7L2*) expression in MSI2-deficient K562 and KU812 cells (Figure 6F and G). Collectively these data indicate that MSI2 can influence Wnt signalling activity through modulation of the Wnt effector LEF-1.

### MSI2 binds LEF1 mRNA and influences its stability

Given the role of MSI2 as an RBP and regulator of post-transcriptional gene expression,^44^ we next assessed whether *LEF1* could be an mRNA partner for MSI2. After first confirming efficient MSI2 RIP in K562 cells (Figure 7A), and enrichment of known mRNA partners *MYB* and *MYC*,^45^ we observed enriched *LEF1* (and *TCF7L2*) mRNA versus control IgG RIP (Figure 7B). Similarly, using an MSI2 CLIP approach we observed enrichment of *LEF1* transcript (and *MYC* and *MYB*) versus IgG CLIP (Figure 7C) indicating that *LEF1* could be a direct mRNA binding target of MSI2. Given β-catenin immunoprecipitates with *LEF1* mRNA through both RIP-seq and RIP-qRT-PCR (Figure 2) but shows limited capacity for direct RNA binding (Figure 3), we hypothesised that β-catenin may impact binding of canonical RBPs like MSI2 to *LEF1* mRNA. To interrogate this, we generated β-catenin depleted K562 cells using two unique shRNAs and re-assessed MSI2 binding to *LEF1* mRNA using MSI2 RIP-RT-qPCR and MSI2-CLIP-RT-qPCR. Firstly, we confirmed β-catenin depletion had no impact on total MSI2 protein (Figure 7D), or overall *LEF1* mRNA expression in β-catenin depleted K562 RIP (Figure 7E) and CLIP inputs (Figure 7F). However, we observed attenuation of *LEF1* mRNA binding with MSI2 upon β-catenin knockdown through both MSI2-RIP (Figure 7G) and MSI2-CLIP (Figure 7H).

**Figure 7.**
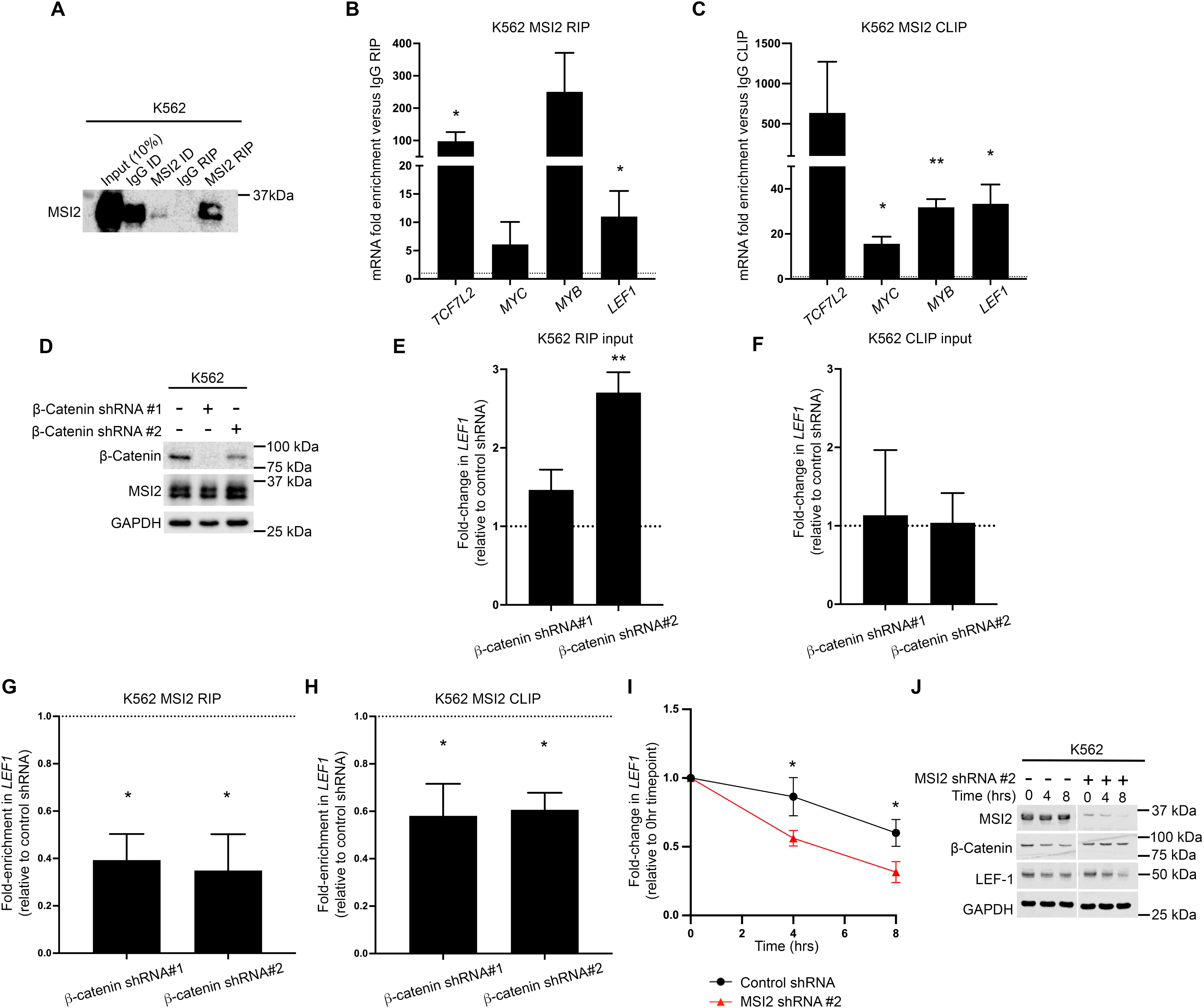
MSI2 regulates LEF-1 expression and binds *LEF1* in a partly β-catenin dependent fashion. **A**) Immunoblot showing MSI2 level in MSI2 RIP performed from K562 cells. ID= immunodepleted lysate. Summary graphs showing the fold enrichment of *TCF7L2*, *MYC*, *MYB*, and *LEF1* mRNA in MSI2 **B)** RIP and **C)** CLIP performed from K562 cells. Fold enrichment is relative to matched IgG RIPs (dashed black line). **D**) Immunoblots showing MSI2 and β-catenin level in K562 cell lines harbouring β-catenin shRNAs versus non-targeting shRNA control. GAPDH indicates protein loading. **E**) Summary graphs showing the fold enrichment of *LEF1* mRNA (versus IgG RIP) in MSI2 RIPs generated from K562 cells +/- β-catenin shRNAs. **G**) Summary graphs showing the fold enrichment of *LEF1* mRNA (versus IgG RIP) in MSI2 CLIPs generated from K562 cells +/- β-catenin shRNAs. **H**) Summary graph showing the fold change in *LEF1* mRNA expression in input K562 cells harbouring β-catenin shRNA versus non-targeting shRNA control (dashed black line) used for MSI2 RIP. **I**) Summary graph showing the fold change in *LEF1* mRNA expression in input K562 cells harbouring β-catenin shRNA versus non-targeting shRNA control (dashed black line) used for MSI2 CLIP. Data represents mean ± 1 s.d (*n* == immunodepleted lysate. Data represents mean3). Statistical analysis is denoted by \**p*<0.05 and \*\**p*<0.01 as deduced from a student’s t-test.

Finally, given MSI2’s prominent roles in post-transcription we assessed *LEF1* mRNA stability following the global inhibition of transcription by actinomycin D (ActD) treatment in K562 cells. Using the K562 variant exhibiting the most complete MSI2 knockdown (shRNA#2) *LEF1* mRNA demonstrated decreased stability compared with the non-targeting shRNA (relative to their respective 0hr timepoint levels; Figure 7I). This corresponded to a markedly accelerated depletion of LEF-1 protein at 8hrs ActD treatment compared with non-targeting shRNA controls (Figure 7J), which is particularly notable given the normally stable half-life we typically observe for the LEF-1 peptide (Supplemental Figure S9). In summary, these data suggest that β-catenin promotes MSI2 binding to *LEF1*, and MSI2 can modulate the subsequent stability of *LEF1* mRNA through a post-transcriptional mechanism.

### MSI2 regulation of LEF-1 may represent an important regulatory axis in human HSPC growth and survival

Both MSI2 and LEF1 have individually documented roles in the maintenance and function of HSPC.^46–50^ Previously, MSI2 has been shown to promote HSPC expansion and we hypothesized that some of this activity could be mediated through LEF-1. To interrogate this hypothesis human CB-derived CD34^+^ HSPC were isolated and lentivirally transduced with ectopic MSI2 expression (Figure 8A) alongside either non-targeting, or LEF-1 shRNAs (Figure 8B). From day 4 onwards (3 days post-lentiviral transduction) we observed a modest but consistent increase in the expansion rates of HSPCs expressing ectopic MSI2 versus control cultures, which was higher (32%±2 increase) at day 8 of liquid culture (Figure 8C) in keeping with previous reports of MSI2-mediated HSPC expansion.^50^ A higher percentage of CD34^+^ events were also observed in MSI2 overexpressing cultures versus controls at this timepoint (Figure 8D). Increases in the expansion rates and CD34 positivity of MSI2 overexpressing cultures were curtailed in the presence of LEF-1 shRNA (Figure 8CD). Primary HSPC cultures harbouring LEF-1 shRNA exhibited consistently reduced growth rates which were lower with shRNA#2 at days 8, 11 and 13 of liquid culture (Figure 8D). Indeed, the growth repression mediated by LEF-1 extended beyond suppressing MSI2-induced expansion, implying independently important roles for LEF1 in human HSPC biology. These data are consistent with our previous study showing LEF-1 can impact the proliferation of myeloid leukaemia cells,^8^ and implies that part of MSI2’s capacity to expand human HSPCs could be mediated through LEF-1.

**Figure 8.**
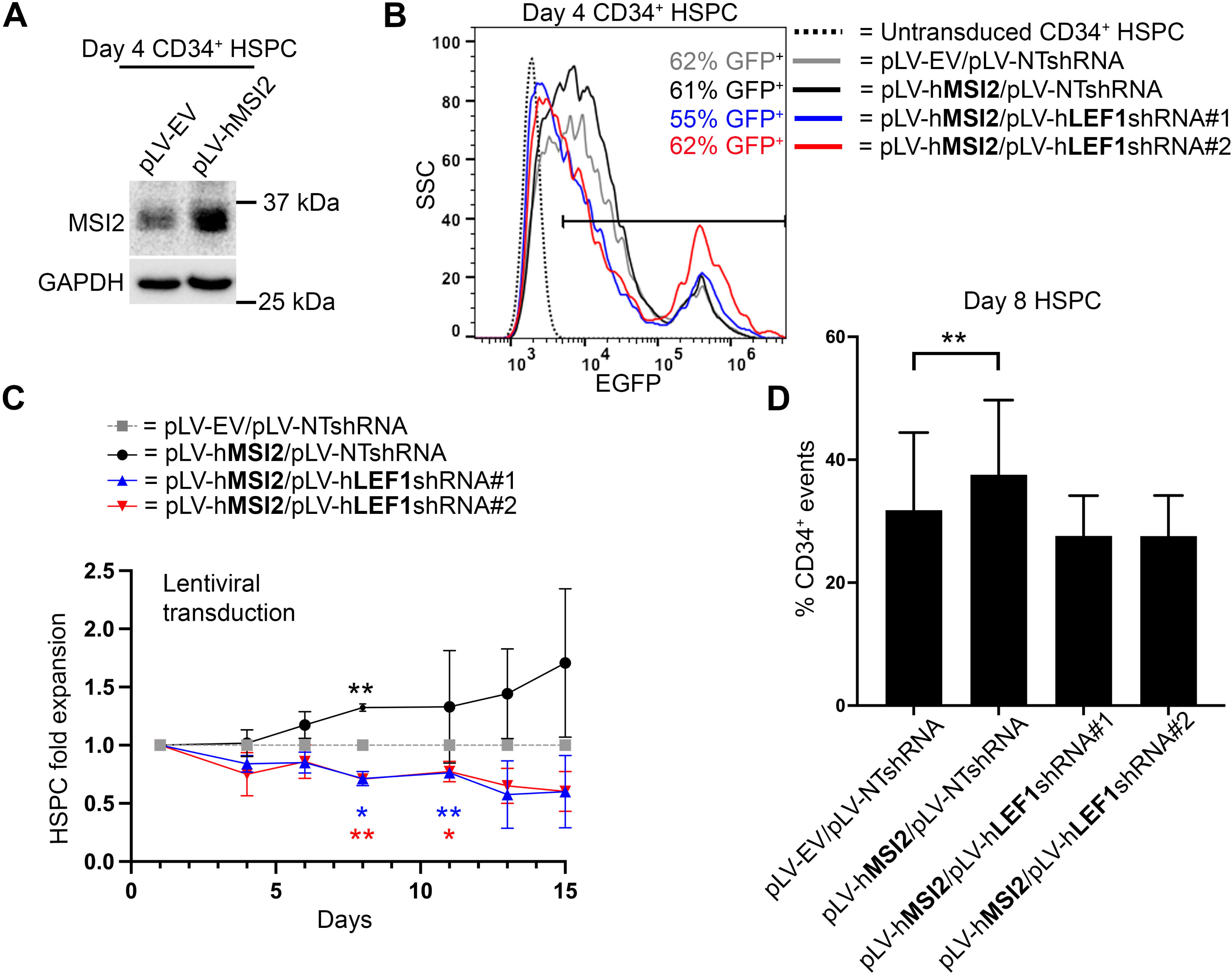
MSI2 mediated human HSPC expansion is partly mediated through LEF-1. **A)** Immunoblot showing the level of MSI2 in day 4 primary CB-derived HSPC cultures following 3 days post lentiviral transduction with empty vector (EV) or human MSI2. GAPDH indicates protein loading. **B**) Representative day 4 flow cytometric histograms plots showing percentage GFP^+^ events in primary HSPC cultures lentivirally transduced with ectopic MSI2 +/- LEF-1 or non-targeting (NT) shRNA, versus matched untransduced cells. **C**) Line graph showing fold-expansion of HSPC cultures following lentiviral transduction with ectopic MSI2 +/- LEF-1 shRNAs. **D**) Bar graph showing the relative percentage of CD34^+^ events in day 8 HSPC liquid cultures following lentiviral transduction with ectopic MSI2 +/- LEF-1 shRNAs. All data represents mean ± 1 s.d (*n* = 3). Statistical analysis is denoted by \**p*<0.05 and \*\**p*<0.01 as deduced from a student’s t-test.

## Discussion

Wnt signalling is a critical pathway regulating human HSCs and is frequently implicated in haematological malignancy. The central mediator of the pathway, β-catenin, has been a longstanding therapeutic target in leukaemia but attempts to therapeutically target the molecule have achieved limited success to date – curtailed by a poor understanding of its molecular interactions in haematopoietic cells. In solid tissues, β-catenin has a vital role in cell adhesion within membrane complexes where it facilitates tight cell-cell adhesion, however such complexes are not a prominent feature of blood cells meaning free β-catenin likely serves alternative functions in this context. Following on from our previous studies showing that β-catenin interacts with vast RBP networks in myeloid cells,^8,9^ we now demonstrate for the first time that β-catenin may contribute towards post-transcriptional regulation in haematopoietic cells.

To identify transcripts over which β-catenin may have post-transcriptional influence in myeloid cells we initially performed β-catenin RIPseq after first confirming β-catenin pulled down with RNA. We identified hundreds enriched transcripts with β-catenin under basal versus Wnt signalling activated conditions across K562 and HEL cells. From K562 cells we identified Wnt/β-catenin signalling as the top signalling pathway enriched from β-catenin associated mRNAs, with many also observed in HEL β-catenin RIPseq. The existence of feedback loops within Wnt signalling has been known for some time with both positive and negative regulators known to be transcriptional targets of the pathway such as *LEF1*,^51^ *LGR5*,^52^ *FZD7*,^53^ and *AXIN2*,^54^ *RNF43*,^55^ *DKK1*,^56^ respectively. However, these were thought to be regulated in a predominantly transcriptional fashion. Our data raise the prospect that β-catenin may also influence feedback loops in Wnt signalling through post-transcriptional mechanisms including both established Wnt target genes, but also Wnt signalling mRNAs that are not transcriptional targets of the pathway including *AMER1*, *BCL9L*, *DVL2* and *CSNK1E*.

The concept that β-catenin could post-transcriptionally govern Wnt signalling is gaining traction with a recent report showing that is it recruited to the translation initiation complex in the dentate gyrus of mice.^57^ Patil and colleagues demonstrated that following eIF4E phosphorylation, β-catenin was substantially recruited to the eIF4E cap complex following long-term potentiation where it preferentially augmented the translation of Wnt signalling mRNAs including *Wnt4*, *Lrp5*, *Fzd2*, *Fzd4* and *Dvl2*. This follows from another study reporting β-catenin recruitment to the messenger ribonucleoprotein and translational pre-initiation complex through the fragile X mental retardation protein (FMRP), where it repressed translation.^58^ Such a post-transcriptional role for free cytoplasmic/nuclear β-catenin might be more prominent in a fluid tissue like blood where its other well characterised role in cell adhesion may not be so relevant. However, trying to disentangle β-catenin’s transcriptional influences from its post-transcriptional effects is technically challenging, especially since data from T-cell acute lymphoblastic leukaemia (T-ALL) cells indicate β-catenin-dependent transcriptional signatures include genes involved with RNA biosynthesis and processing.^59^

The capacity for β-catenin to bind RNA directly remains unclear. Previously, β-catenin has been shown to bind selective mRNAs including COX2 through the 3’UTR AU-rich elements in concert with RBPs such as HuR,^38,42^ and recently has been shown to bind double stranded RNA.^43^ Numerous reports also exist of β-catenin binding and modulating the stability/splicing of numerous transcripts including *CA9*, *SNAI1*, *IL6*,^60^ *Cadherin 11*,^61^ and an estrogen receptor-β (*ER-*β) variant.^62^ Armadillo domains have also been identified as one of the most frequent domain types associated with RNA binding amongst non-canonical RBPs,^41^ and β-catenin has 12 such repeating structures in its central domain. However, β-catenin has no recognised canonical RNA binding motif, and we observed little RNA pull down through a more direct CLIP assessment in myeloid cells. Therefore, we hypothesise that β-catenin’s interactions with RNA in this context occur indirectly through binding other canonical RBPs and there were many identified in our original β-catenin interactome study.^8,9^ Thus far, we have confirmed interactions with RBPs such as WT1,^27^ HuR (Supplemental Figure S1A), LIN28B and TOE1 (unpublished) and all of these β-catenin:RBP interactions occur irrespective of RNA presence (Figure 4BC), suggesting β-catenin complexes with RBPs, rather than associates though co-occupancy of RNA. This follows from many other reported interactions of β-catenin with RBPs including FUS,^63^ HuR,^38,60^ and FMRP,^58^ and suggests a wider role for β-catenin in RNA:RBP complexes.

We were the first to report functional crosstalk between Wnt/β-catenin signalling and MSI2 in myeloid cells.^64^ A subsequent report from the Sheng group adopted a deletion of the chromosome 5q (del(5q)) murine model of MDS which has previously demonstrated a dependency on the negative Wnt regulator and tumour suppressor APC for MDS development.^15^ The authors show that MSI2 may also impact Wnt signalling function in murine HSC, however there no established measurements of Wnt signalling activity (e.g. TCF/LEF activity, Wnt target gene assessment, β-catenin stabilisation/nuclear localision dynamics) performed in this study.^65^ We found the principle connection between these two proteins appeared to be the Wnt transcription factor LEF-1 which was suppressed with MSI2 knockdown across multiple cell lines. This aligns with a previous report of lower LEF-1 (and TCF4) levels (protein and mRNA) upon MSI2 depletion in SMMC-7721 hepatocellular carcinoma cells.^66^ A Wnt gene signature was also observed in response to MSI2 loss in four human leukaemia cell lines in a study by Kharas *et al*, which were reciprocally upregulated upon ectopic MSI2 expression in murine HSC.^47^ Given the well documented importance of MSI2 to HSPCs,^46,67,68^ we reasoned that regulation of *LEF1* might be a mechanism utilised by MSI2 to control HSPC growth and survival. Indeed, *LEF1* knockdown perturbed MSI2-mediated expansion of HSPCs, however we cannot rule out *LEF1* acting independently of any MSI2-mediated regulation since it is also a key regulator of the HSPC compartment.^49,69^

Finally, this study partly examined the mechanism by which MSI2 could regulate *LEF1* expression and found that MSI2 bound *LEF1* mRNA through both RIP and CLIP approaches. This is in agreement with an elegant study by the Pillai group who also performed MSI2 iCLIP in K562 cells and we noted that *LEF1* was a high confidence mRNA partner in their dataset.^70^ Interestingly, they found only 2.6% of 4,000 high confidence MSI2 mRNA partners (mainly via 3’-UTR) are subsequently subject to translational regulation by MSI2. Our study also suggests that perhaps mRNA stability, rather than translation, is the key mechanism regulating *LEF1* expression, concordant with MSI2’s previously documented roles in modulating mRNA stability.^71–74^ We observed the degree of MSI2:*LEF1* binding could be impacted by β-catenin level (with no consistent or substantial overall change to total *LEF1* or MSI2 levels) alluding to a wider role for β-catenin augmenting RBP:RNA complexes in myeloid cells. How β-catenin influences RBP:RNA complexes remains unknown and could represent a novel function for β-catenin in haematopoietic cells. In its well characterised transcriptional role it recruits TCF/LEF transcription factors and transcriptional co-activators (CBP, p300) to the DNA promoters of Wnt target genes, and perhaps β-catenin performs similar scaffolding functions for facilitating specific RBP:RNA interactions, particularly those implicating Wnt signalling mRNAs. However, trying to resolve its well characterised transcriptional effects, from any potential post-transcriptional influences, will be technically challenging, especially since both nascent RNA and active RNA polymerase II have been recently been observed within β-catenin condensates.^39^ Further work is now required to define β-catenin’s post-transcriptional versus transcriptional roles in a haematological context.

## Supporting information

Supplementary data file (beta-catenin RIPseq)

## Acknowledgements

This work was funded by the Kay Kendall Leukaemia Fund (KKL1051/KKL1446), Leukaemia & Myeloma Research UK (4-5/06.21R) and the Children’s Cancer & Leukaemia Group (CCLGA 2023 16 Morgan). We are grateful to the Leeds Genomics facility for executing RNA sequencing experiments. Kind thanks to Dr Paraskevi Diamanti (University of Bristol) for AML patient sample collection and Lizzy Hoole (Institute for Child Life & Health, UHBristol & Weston NHS Foundation Trust) for supplying clinical data. Thanks to the midwives and clinical research nurses at University Hospitals Sussex (UHS) NHS trust including Raquel Akieme, Valentina Toska, Lorraine Shah-Goodwin, Denise Skinner, Elohor Uwadiogbu and Caroline Humphreys, for the collection of human umbilical cord blood. We are indebted to the patients and their families who gave consent for their cells to be used for our research.

## Author contributions

MW performed experiments, analysed data, co-wrote the manuscript and managed laboratory. OS completed RIP-seq analyses, AG performed CLIP analyses, MR generated library preps and ran RNA sequencing whilst HP and EM performed AlphaFold predictive modelling. DN provided experimental advice, whilst TC and AB sourced primary clinical samples. LC, SN and BT provided RNA experimental guidance and reagents. RGM performed experiments, analysed data, co-wrote the manuscript, secured funding and directed the study.

