## Supplementary figures for "β-Catenin associates with a Wnt signaling mRNA network in myeloid cells through canonical RBP binding"

**Supplementary Table S1. Clinical characteristics of AML patient diagnostic/relapse samples used in this study**.

| **Patient no.** | **Age (at diagnosis)** | **Sex** | **WBC count (x10^9^/L)** | **Sample type** | **Secondary disease (Y/N)** | **Genetic information** | **Other clinical information** |
| --- | --- | --- | --- | --- | --- | --- | --- |
| 1 | 4 | F | 2.6 | BM | N | Normal karyotype, NPM1^+^ (exon 12), FLT3^-^ | MRD neg. CD13+, CD33+, CD34+, CD117+, MPO+ |
| 2 | n/a | n/a | n/a | n/a | n/a | n/a | n/a |
| 3 | n/a | n/a | n/a | n/a | n/a | n/a | n/a |
| 4 | 14 | F | 7.6 | BM | N | MLL rearrangement. Karyotype: 46,XX,ins(10;11)(q11.2;q23.1q23.3).ish ins(10;11)?inv(11)(q23.3)(5’MLL+)(q23.1)(3’MLL+) | BMT for high-risk AML. CD33+, MPO+, CD34-, CD117+, TdT+, CD64+, CD11c+, CD15+, CD11b+, NG2+ |
| 5 | n/a | n/a | >200 | PB | Y | n/a | Post-allogeneic transplant. M0/1 (previously diagnosed with M3 10 years previous) |
| 6 | 6 | M | 70.7 | n/a | Y | n/a | Secondary to Ewings Sarcoma. Myelomonocytic morphology. Deceased. |
| 7 | 5 | M | 1.6 | BM | N | n/a | M5. Deceased. |
| 8 | 16 | F | 20.6 | BM | N | Normal karyotype 46,XX[20] | CD33+, MPO+, CD34+, CD117+, CD13+, CD14-, CD7+, CD45 weak, CD11c+, TdT- |
| 9 | 10 | F | n/a | BM | n/a | n/a | Deceased |
| 10 | 76 | F | 389 | LP | N | Normal karyotype, NPM1^+^, FLT3^+^ | N/A |
| 11 | 7 | M | 3.5 | BM | N | t(8;21)(q22;q22) RUNX-RUNX1T1  45,X,-Y,t(8;21)(q22;q22)[9]/46,XY[1] | 8% myeloid blasts present (CD13+, CD33+, CD34+, CD117+, MPO+) |
| 12 | 7 | F | n/a | BM | N | MPAL, 46XX, del5q, abnormal 21 | n/a |
| 13 | 64 | F | 13.3 | BM | N | Normal karyotype | AML with underlying MDS like changes |
| 14 | 7mo | F | 168.6 | BM | N | Karyotype: 47,XX,+21[5]/46,XX[5] nuc ish(CBFA2T3,GLIS2)X3(CBFA2T3 con GLIS2x2)[92/150]  FISH showed no evidence of CBFB or RUNX1-RUNX1T1 rearrangement however, an additional copy of RUNX1 was found.  33+, CD34+, CD117+, MPO+, DR-, CD13+ | BMT 24/12/20 due to high risk genetics.  Alive, in remission.  Chromosome and FISH analysis showed an inv(16)(p13.3q24.3) [CBFA2T3-GLIS2] rearrangement and trisomy for chromosome 21. This is consistent with a diagnosis of AML and the CBFA2T3-GLIS2 rearrangement is a poor risk finding according to the MyeChild 01 protocol (received Gemtuzumab as part of Myechild trial treatment.) |
| 15 | 17 | M | n/a | BM | Y | n/a | Post-BMT for AML following 2 relapses. Deceased |
| 16 | 4 | M | 3.2 | BM | N | t(10;11)(p11.2;q23) KMT2A-MLLT10. Cytogenetically cryptic. KMT2A ex8-MLLT10 ex9 or KMT2A ex9-MLLT10 ex10 fusion detected. NPM1^-^FLT3^-^ | CD13-, CD33+, CD34-, CD117+/-, CD11c+, CD64+, CD14-, NG2+. High risk cytogenetics. BMT. MRD neg post course 1+2. |
| 17 | n/a | n/a | n/a | n/a | n/a | n/a | n/a |
| 18 | 15 | F | 6.5 | BM | N | MLL (KMT2A) rearrangement, t(10;11)(p11-p14,q23), MLL-MLLT10 | MRD detected post treatment course 1 |
| 19 | 2 | M | 2.4 | BM | N | t(9;11)(p22;q23), t(11;21)(q23;q8) | n/a |
| 20 | 7 | F | 34.4 | BM | Y | MLL rearrangement t(9;11) | M5a morphology. BMT following relapse. Deceased |

BM = Bone marrow

PB = Peripheral blood

LP = Leukapheresis

MRD = Minimal residual disease

BMT = Bone marrow transplant

AML= Acute myeloid leukemia

MDS= Myelodysplastic syndrome

MPAL= Mixed phenotype acute leukemia

MLL = *Mixed-lineage leukemia*

NPM1 = *Nucleophosmin*

FLT3 = *Fms-like tyrosine kinase 3*

RUNX1 = *Runt-related transcription factor 1*

GATA2 = *GATA Binding Protein 2*

**Supplementary Table S2 - Forward and reverse primer sequences used for RT-qPCR.**

| **Primer** | **5’ -> 3’ sequence** |
| --- | --- |
| *ACTB* Fw | TTGTTACAGGAAGTCCCTTGCC |
| *ACTB* Rv | ATGCTATCACCTCCCCTGTGTG |
| *GAPDH* Fw | ACAGTCAGCCGCATCTTCTT |
| *GAPDH* Rv | ACGACCAAATCCGTTGACTC |
| *AMER1* Fw | AGTACCCGTGAACAAAGAGCA |
| *AMER1* Rv | AGGCAGTACAGATACCCTTC |
| *BCL9L* Fw | TGAACCTGAACGTGCAGATGA |
| *BCL9L* Rv | CCCTGGTTGGGAAACTGTG |
| *AXIN2* Fw | TTGGCTACTCCGTAAAGTTTTGGT |
| *AXIN2* Rv | TACACTCCTTATTGGGCGATCA |
| *LEF1* Fw | AGAACACCCCGATGACGGA |
| *LEF1* Rv | GGCATCATTATGTACCCGGAAT |
| *TCF7L2* Fw | AGAAACGAATCAAAACAGCTCCT |
| *TCF7L2* Rv | CGGGATTTGTCTCGGAAACTT |
| *MYB* Fw | GAAAGCGTCACTTGGGGAAAA |
| *MYB* Rv | TGTTCGATTCGGGAGATAATTGG |
| *MYC* Fw | AGCGACTCTGAGGAGGAA |
| *MYC* Rv | CCAGCAGAAGGTGATCCA |
| *RNA18SN1* Fw | CTCAACACGGGAAACCTCAC |
| *RNA18SN1* Rv | CGCTCCACCAACTAAGAACG |

**Supplementary Table S3 – Lentiviral plasmids used in this study for transgene expression.**

| **Target gene** | **Expression type** | **Vector type** | **Supplier** |
| --- | --- | --- | --- |
| Non-targeting | shRNA control | pLKO.1-puro Non-Mammalian shRNA Control Plasmid  SHC002 | Merck MISSON® |
| *MSI2* | shRNA (#1) | pLKO_TRCN0000062808 | Merck MISSON® |
| *MSI2* | shRNA (#2) | pLKO_TRCN0000062809 | Merck MISSON® |
| *CTNNB1* | shRNA (#1) | pLKO_TRCN0000314920 | Merck MISSON® |
| *CTNNB1* | shRNA (#2) | pLKO_TRCN0000314921 | Merck MISSON® |
| Empty vector control | Ectopic | pLV[Exp]-Puro-EF1A>ORF_91bp  (VB160723-1006snj) | VectorBuilder |
| *MSI2* | Ectopic | pLV[Exp]-Puro-EF1A>hMSI2[NM_138962.4]  Vector ID:VB230612-1123ebn | VectorBuilder |
| Empty vector control | Ectopic | pLV[Exp]-mCherry-EF1A>ORF_91bp  (VB230711-1149jnd) | VectorBuilder |
| *MSI2* | Ectopic | pLV[Exp]-mCherry-EF1A>hMSI2[NM_138962.4]  Vector ID:VB230612-1125btf | VectorBuilder |
| Non-targeting | shRNA control | pLV[shRNA]-EGFP-U6>Scramble[shRNA#2] (Vector ID:VB230321-1431mhe) | VectorBuilder |
| *LEF1* | shRNA (#1) | pLV[shRNA]-EGFP-U6>hLEF1  TRCN0000418104 | VectorBuilder |
| *LEF1* | shRNA (#2) | pLV[shRNA]-EGFP-U6>hLEF1  TRCN0000428355 | VectorBuilder |

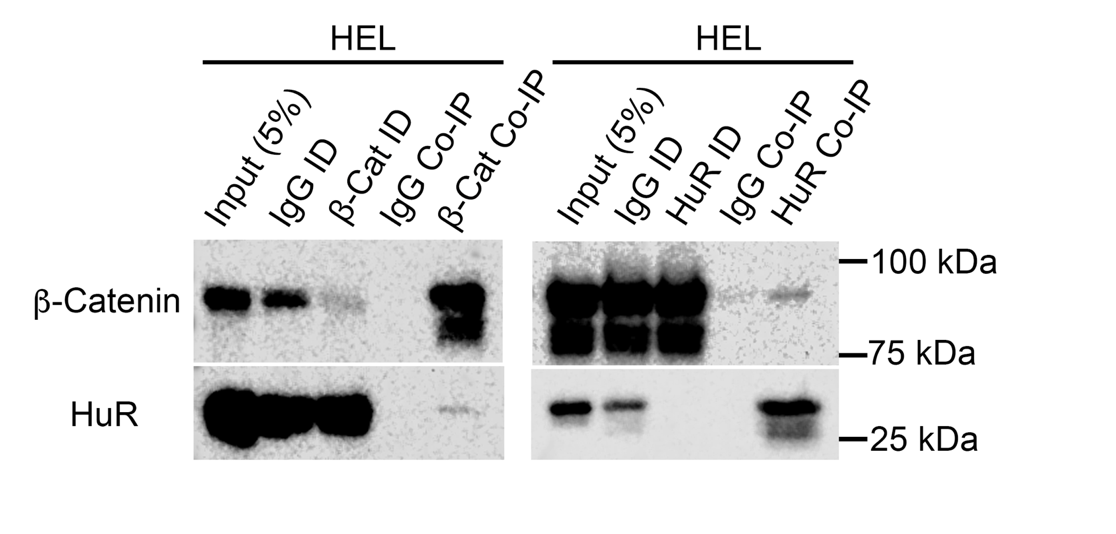

**Supplementary Figure S1. HuR and β-Catenin interact in AML cells.** Immunoblots showing the level of HuR (*ELAVL1*) protein present in β-catenin Co-IP, and reciprocally the level of β-catenin present in HuR Co-IP from whole cell lysates derived from HEL cells.

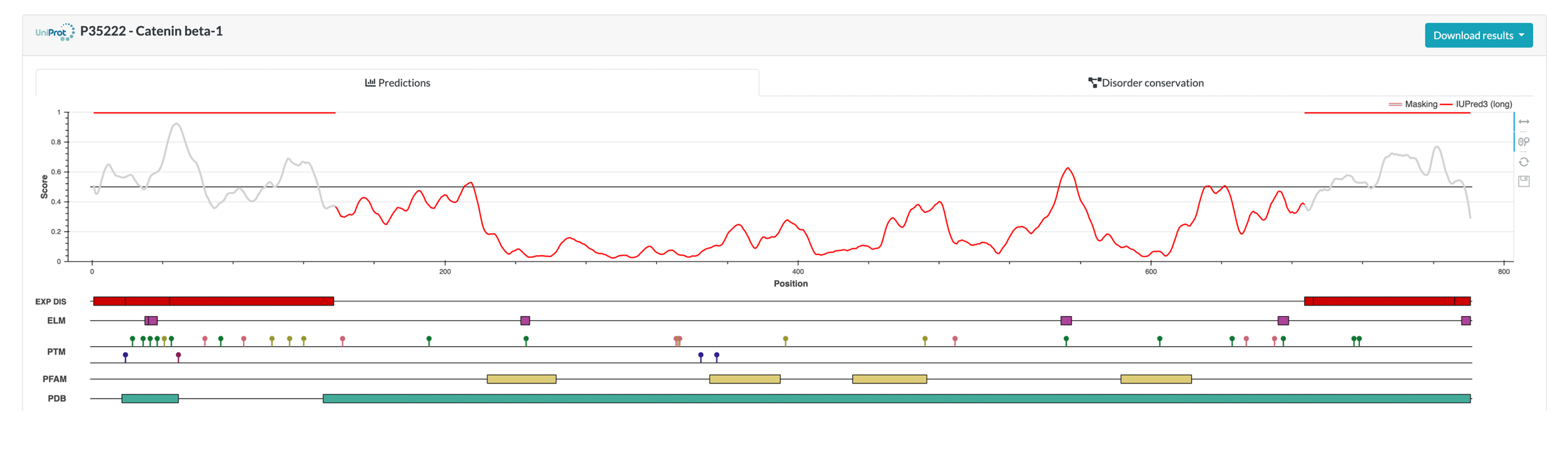

**Supplementary Figure S2. β-Catenin intrinsically disordered regions (IDRs).** Amino acid residue map from IUPred3 server predicting the most likely location within the β-catenin peptide for IDRs which are the N- and C-termini with values between high confidence values between 0.5-1 (Erdos, G. et al. *Nuc Acids Res*, 2021).

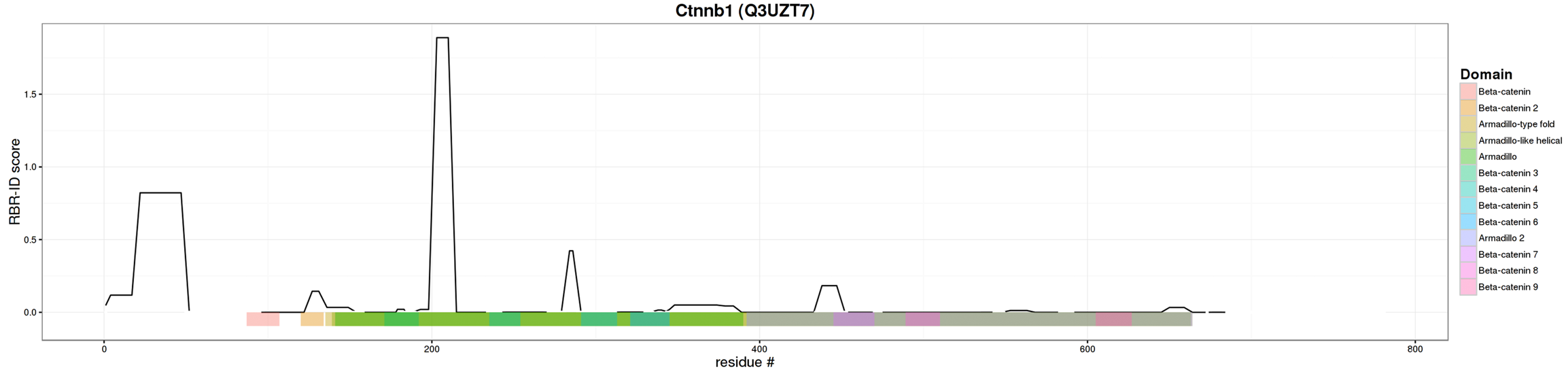

**Supplementary Figure S3. β-Catenin central armadillo domains predict RNA binding.** Amino acid residue map from Bonasio Lab RBR-ID browser v0.3 showing likely RNA-binding regions (RBR) in the ARM domain (seq: TMQNTNDVETAR, RBR-ID score: 1.89) and N-terminus (seq:

AAVSHWQQQSYLDSGIHSGATTTAPSLSGK, RBR-ID score: 0.82) of β-catenin from nuclei of murine embryonic stem cells following RBP screen (He, C. *et al*, *Mol Cell*, 2016).

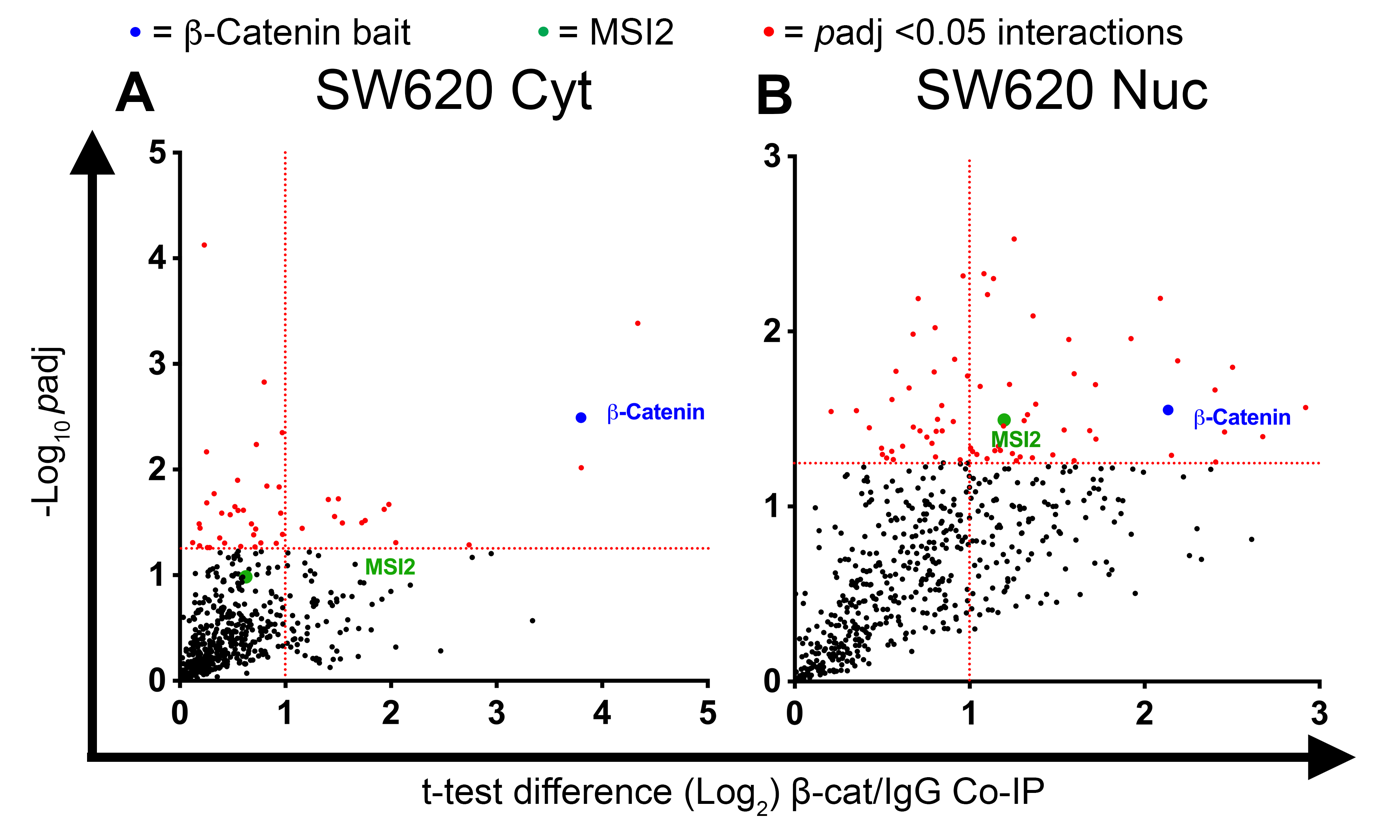

**Supplementary Figure S4. Proteomics analyses reveal β-catenin:MSI2 interaction in colorectal cancer cells.** Scatter plots showing β-catenin protein interactions detected in SW620 cytosolic and nuclear fractions. Vertical dashed red line indicates the threshold for 2-fold change in protein binding at log_2_ (=1) relative to IgG co-immunoprecipitation. Horizontal red line represents *P*=0.05 on log_10_ scale (=1.3). Highlighted red dots indicate interactions where p<0.05. Remaining black dots represent other proteins detected in the MS analysis, green dot highlights MSI2 detection. Fold change values less than 0 are not shown because these likely represent contaminants.

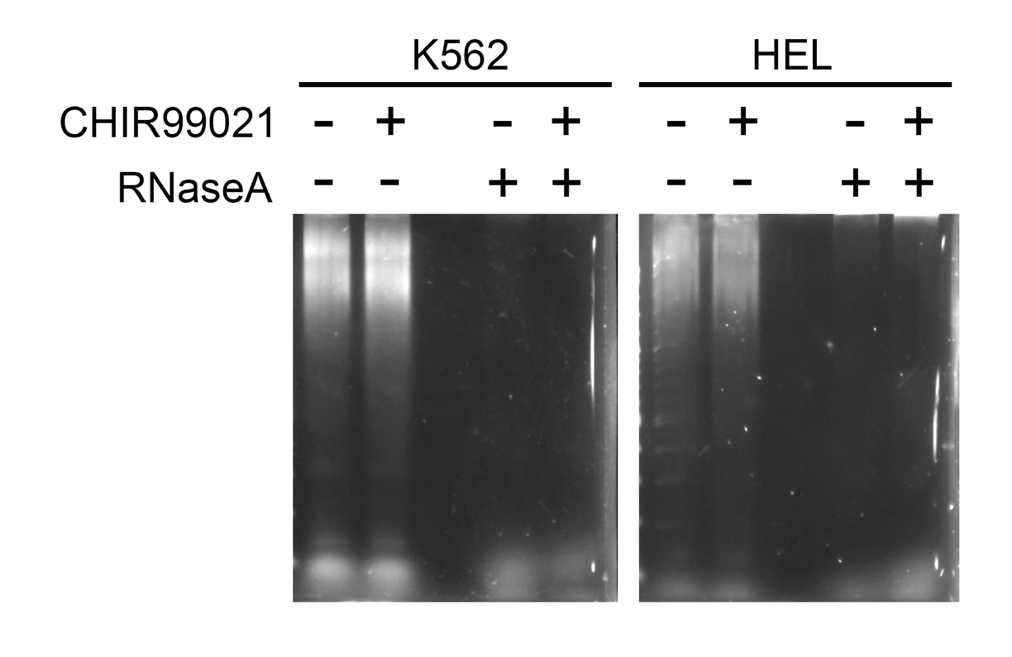

**Supplementary Figure S5. RNA digestion through RNaseA treatment.** Agarose gel electrophoresis showing the stability of total RNA in K562 and HEL lysates treated overnight +/-5mM CHIR99021 (or respective DMSO control) and +/- 20 mg/mL RNaseA.

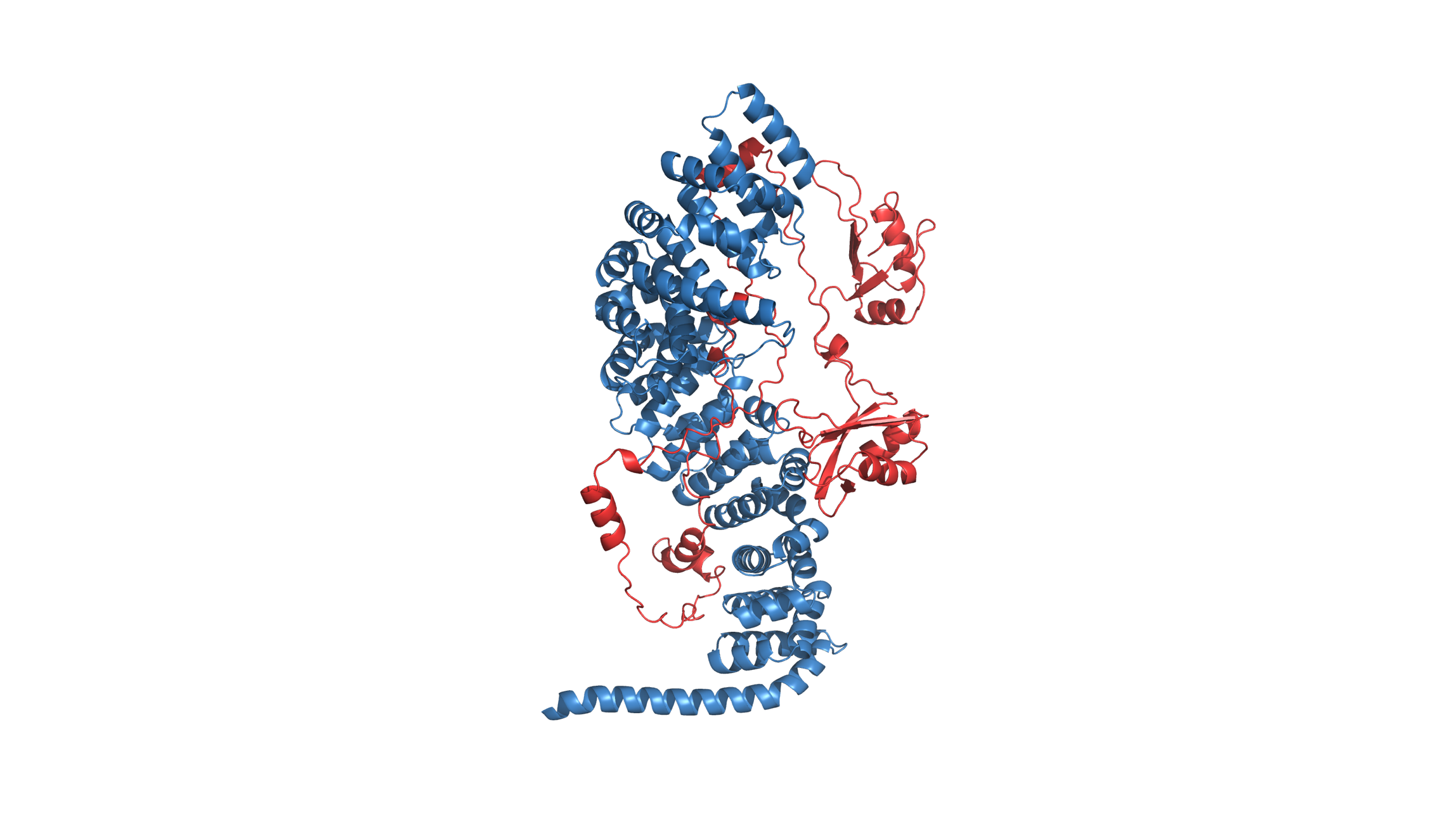

**Supplementary Figure S6. AlphaFold 3 modelling of the β-catenin:MSI2 interaction.** The β-catenin molecule is shown in blue, whilst the MSI2 molecule is shown in red. Utilising AlphaFold 3, we generated predictions for the structure of the β-catenin:MSI2 interaction. Even in the model with the highest confidence metrics, the predicted template modelling (pTM) and Interface pTM (ipTM) scores were low (0.54 and 0.29 respectively). To further investigate the predicted interaction, we used PDBePISA to analyse the protein-protein interface (Krissinel, E and Henrick K, *J. Mol. Biol*, 2007). Despite a sizeable buried hydrophobic interface, the complexation significance score of 0 suggested that the interface is not significant for assembly formation. Overall, the analysis suggests that the nature of the β-catenin:MSI2 interaction may be indirect, possibly as part of a larger complex (Jumper J, et al. *Nature*, 2021; Evans R, et al. *bioRxiv*, 2022).

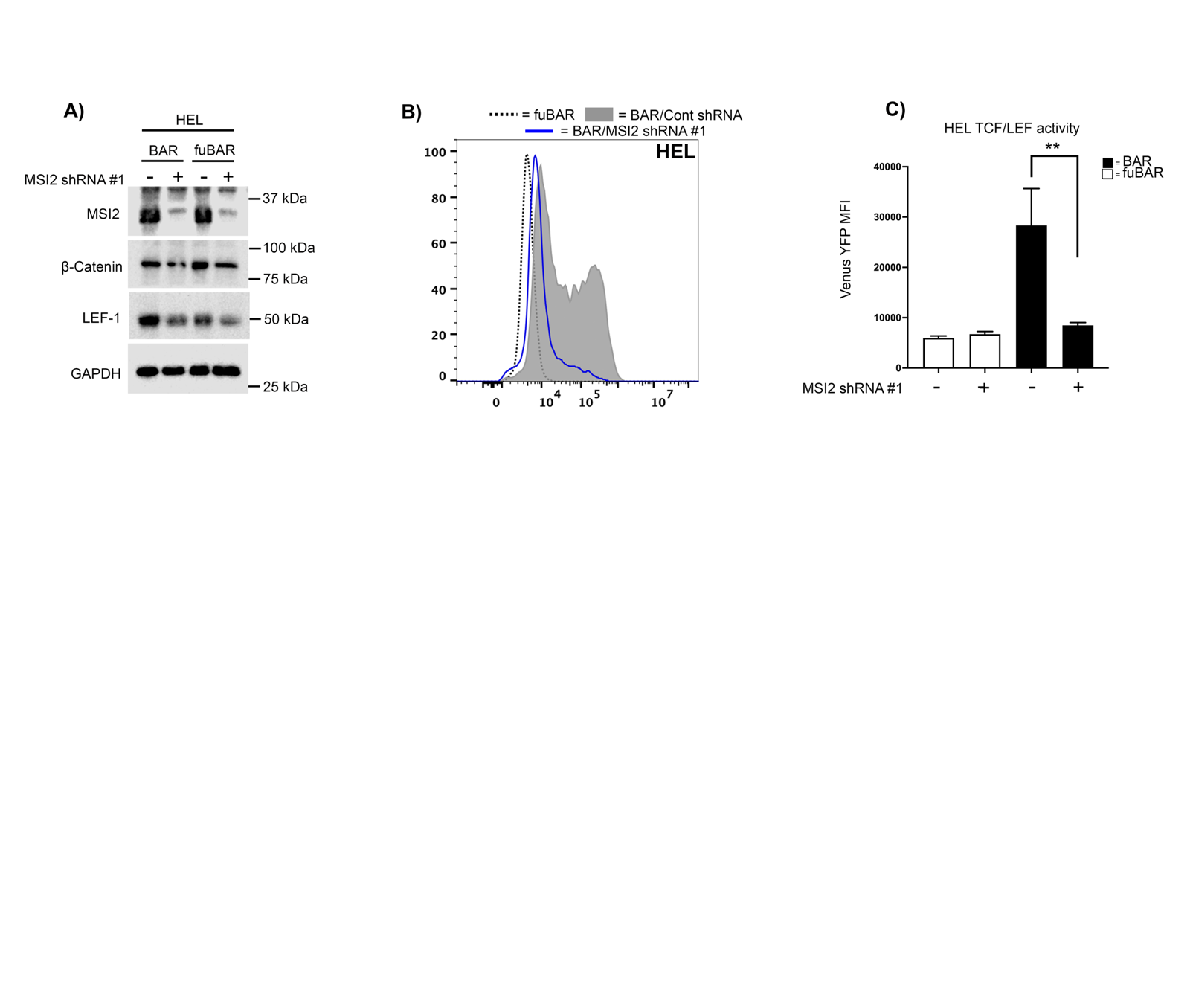

**Supplementary Figure S7. MSI2 knockdown impairs Wnt signalling output in HEL cells. A)** Immunoblots showing total MSI2, β-catenin and LEF-1 level in HEL cells (Wnt reporter BAR and fuBAR variants) harbouring MSI2 shRNA or non-targeting shRNA control. GAPDH indicates protein loading. **B)** Representative flow cytometric histograms showing YFP intensity BAR or negative control fuBAR HEL cells +/- MSI2 shRNA following treatment with 5mM CHIR99021 overnight. The fuBAR (dashed), non-targeting control shRNA (grey filled), and MSI2 shRNA (blue) histograms are shown. **C)** Summary bar graphs showing the median YFP fluorescence intensity (MFI) generated from the BAR/fuBAR in HEL +/- MSI2 shRNA with +/- 5mM CHIR99021. All data represents mean ± 1 s.d (*n* = 3). Statistical analysis is denoted by ***p*<0.005 as deduced by a student’s paired t-test.

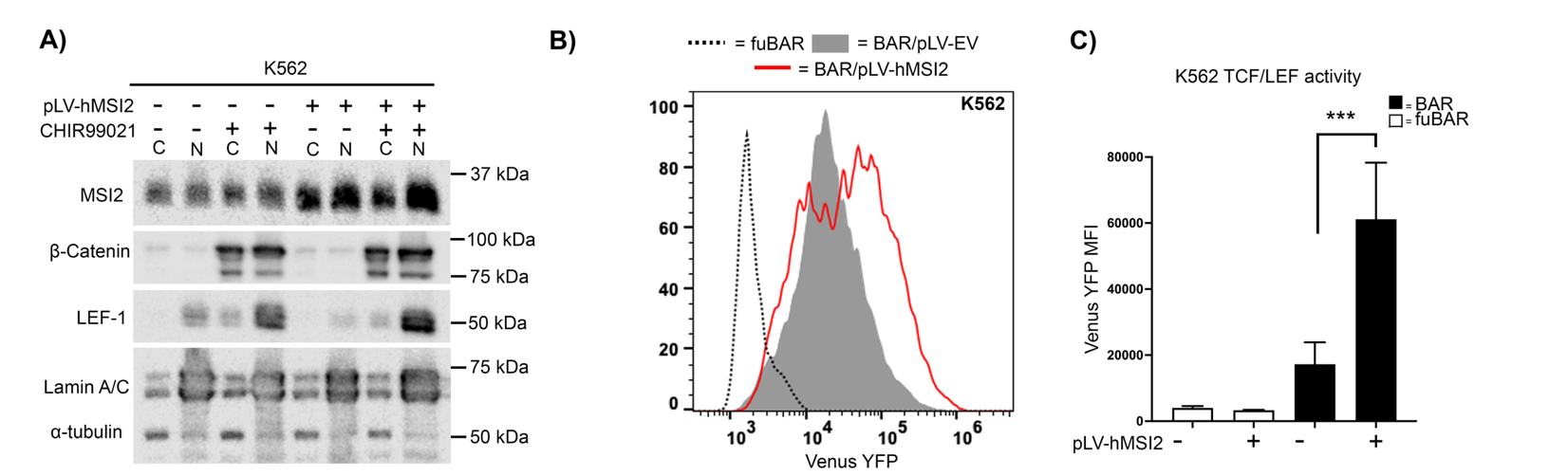

**Supplementary Figure S8. MSI2 overexpression enhances Wnt signalling in K562 cells. A)** Immunoblots showing total β-catenin, LEF-1 and MSI2 subcellular localization in K562 cells lentivirally transduced with ectopic human MSI2 (pLV-hMSI2) +/- 5mM CHIR99021. Lamin A/C and α-tubulin indicate the purity/loading of the nuclear (N) and cytosol (C) fractions, respectively. **B)** Representative flow cytometric histograms showing intensity of the TCF-dependent expression of Venus YFP from the BAR reporter, or negative fuBAR control in K562 cells +/- ectopic MSI2 OE shRNA following treatment with 5mM CHIR99021 overnight. The fuBAR (dashed), non-targeting control shRNA (grey filled), and MSI2 OE shRNA (red) histograms are shown. **C)** Summary bar graphs showing the median fluorescence intensity (MFI) generated from the BAR/fuBAR in K562 +/- MSI2 OE shRNA with +/- 5mM CHIR99021. All data represents mean ± 1 s.d (*n* = 3). Statistical anlaysis is denoted by ****p*<0.0005 and as deduced by a paired t-test.

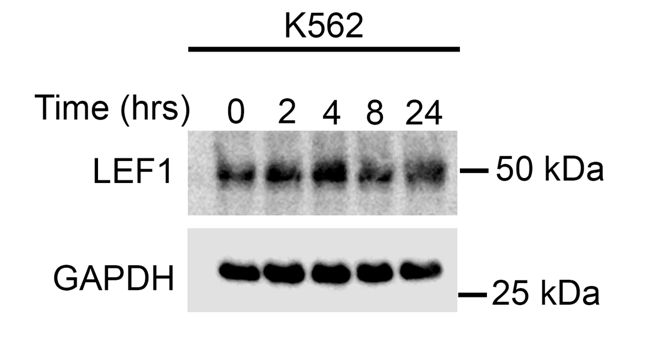

**Supplementary Figure S9. The LEF-1 peptide has a long half-life.** Immunoblot showing the total level of LEF-1 protein following treatment with 5μg/ml of Actinomycin D exposure for 2, 4, 8 and 24 hours. GAPDH is used to indicate protein loading.
